# Novel populations of CD4^+^ T cells associated with vaccine efficacy

**DOI:** 10.1101/2022.06.23.497400

**Authors:** Therese Woodring, Colin N. Dewey, Lucas Dos Santos Dias, Xin He, Hannah E. Dobson, Marcel Wüthrich, Bruce Klein

## Abstract

Memory T cells underpin vaccine-induced immunity but are not yet fully understood. To distinguish features of memory cells that confer protective immunity, we used single cell transcriptome analysis to compare antigen-specific CD4^+^ T cells recalled to lungs of mice that received a protective or nonprotective subunit vaccine followed by challenge with a fungal pathogen. We unexpectedly found populations specific to protection that expressed a strong type I interferon response signature, whose distinctive transcriptional signature appeared unconventionally dependent on IFN-γ receptor. We also detected a unique population enriched in protection that highly expressed the gene for the natural killer cell marker NKG7. Lastly, we detected differences in TCR gene use and in Th1- and Th17-skewed responses after protective and nonprotective vaccine, respectively, reflecting heterogeneous *Ifng*- and *Il17a-*expressing populations. Our findings highlight key features of transcriptionally diverse and distinctive antigen-specific T cells associated with protective vaccine-induced immunity.

## INTRODUCTION

Vaccines have saved millions of lives, eradicated fatal diseases, and proved essential to controlling emerging infectious disease threats (Duclos et al., 2009; Galvani et al., 2021; Heaton, 2020). While vaccines were initially developed without mechanistic understanding of immunity, they are now recognized to require the response of antigen-specific T cells that produce cytokines to activate phagocytes, induce antibody production in memory B cells, and persist long after the initial antigen challenge. These memory T cells—including circulating effector memory T cells (T_EM_), central memory T cells (T_CM_), and tissue-resident memory cells (T_RM_)—may be elicited by a variety of vaccine types and confer variable protection based on the route of vaccination and magnitude of the initial T cell response (Panagioti et al., 2018; Pollard and Bijker, 2021; Schenkel and Masopust, 2014). In contrast to circulating antibodies, however, memory T cells can be difficult to isolate from the periphery and remain poorly characterized. Given ongoing challenges in developing vaccines that induce cellular immunity against some of the most important global pathogens, a better understanding of these T cells—and what distinguishes protective from nonprotective vaccine-induced T cell responses—is a priority.

Single cell transcriptome analysis (scRNAseq) is one tool for characterizing memory T cells at the site of pathogen encounter. Whereas traditional methods classify cells by expression of a small subset of known markers, scRNAseq defines cell phenotypes agnostically, by gene expression profiles across the entire transcriptome. Often, the approach validates known differences in cell types; however, it may also expand or even challenge traditional frameworks for classifying complex populations. For instance, scRNAseq analysis groups T cells from blood, lymphoid, and lung tissue by activation states distinct to known CD4^+^ and CD8^+^ lineages, whereas effector T helper cells responding to various colonic pathogens do not segregate into canonical Th1, Th2, and Th17 archetypes (Kiner et al., 2021; Szabo et al., 2019).

Sequencing-based methods also identify novel cell types (Stubbington et al., 2017). One such cell, observed in recent scRNAseq experiments, is the type I interferon-signature T cell (Andreatta et al., 2021; Arazi et al., 2019; Gowthaman et al., 2019; Harsha Krovi et al., 2020; Kiner *et al*., 2021; Seumois et al., 2020; Singhania et al., 2019; Szabo *et al*., 2019; Tibbitt et al., 2019; Zemmour et al., 2018). These cells (hereafter “Tis T cells”) are distinct for the striking upregulation of multiple genes that typically are induced by type I interferons (IFN) and have well established roles in cellular responses to viral infection. However, these Tis T cells have appeared across diverse immunological settings where type I interferon would not be expected, such as dust mite allergy, *Alternaria* sensitization, and *Salmonella* and *Citrobacter* infection (Gowthaman *et al*., 2019; Kiner *et al*., 2021; Tibbitt *et al*., 2019). Their function remains unknown.

Herein, we compare the transcriptional phenotypes of antigen-specific CD4^+^ T cells recalled to lungs of mice challenged with lethal pulmonary fungal infection after they received a subunit vaccine that is highly protective when given subcutaneously (SC), but not intranasally (IN). By using single cell transcriptome analysis, we uncover populations of T cells previously unrecognized in the setting of vaccine induced protective immunity. For example, we uncovered two T cell populations that express a strong type I interferon response signature (Tis), unexpected in the context of antifungal immunity, but consistent with descriptions of the novel Tis T cell phenotype recently reported in this journal. Unique to our report, we observe increased abundance of Tis T cells only during a protective immune response, together with the unconventional dependence of the Tis signature on IFN-γ receptor. We also highlight a unique CD4^+^ T cell population enriched in protection that bears many NK cell markers including *Nkg7*, of recent interest for its regulatory role in CD4^+^ T cell activation and pathogen control. Finally, while we validate previously described Th1- and Th17-skewed responses after protective and nonprotective vaccination, respectively, we uncover features of Th1 and Th17 responses that reflects a tension between the widely accepted framework of conventional T helper cell archetypes (Th1, Th2, Th17) and the nuance that can be detected by newer, hypothesis-free approaches to immune cell profiling

## RESULTS

### scRNAseq analysis of antigen-specific memory CD4^+^ T cells from intranasally (IN) and subcutaneously (SC) vaccinated, *Blastomyces*-challenged mice

Mice were vaccinated either intranasally (IN) or subcutaneously (SC) with *Blastomyces dermatitidis* endoglucanase-2 (Bl-Eng2) six weeks prior to pathogen challenge (Fig. 1a). As described previously, these routes of vaccine delivery both induce substantial numbers of antigen-specific T cells but are associated with divergent outcomes in response to lethal experimental challenge with *B. dermatitidis*. Mice vaccinated SC effectively control lung fungal burden, whereas mice vaccinated IN do not (Dobson et al., 2020). For scRNAseq analysis, tetramer-positive CD4^+^ T cells were FACS sorted and sequenced 3 days after the pulmonary challenge with *B. dermatitidis* (Fig. 1b; Supplemental Fig. 1).

**Figure 1.**
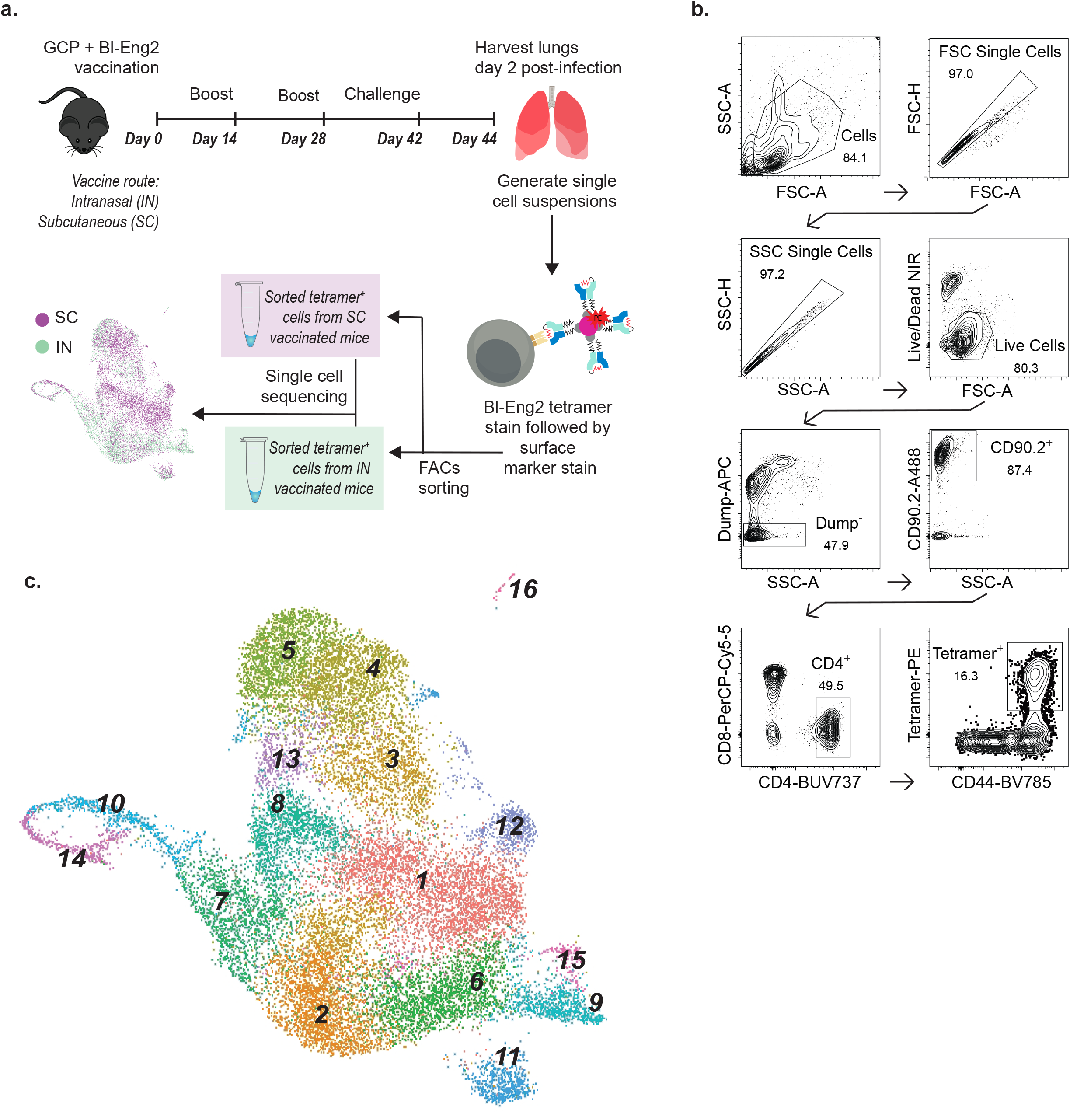
Single-cell RNAseq analysis of tetramer-positive T cells in *Blastomyces-*challenged mice following subcutaneous (SC) or intranasal (IN) vaccination. **(a)** Experimental schema for IN and SC vaccination with *Blastomyces* endoglucanase-2 (Bl-Eng2), lethal experimental challenge with *Blastomyces*, sorting of Bl-Eng2-specific CD4^+^ T cells with tetramer, and single-cell RNAseq. **(b)** Gating strategy for selection of tetramer^+^ CD4^+^ T cells. Representative flow cytometry plots shown for the SC group cells; see Supplemental Figure 1 for IN group flow cytometry plots. **(c)** Uniform manifold approximation and projection (UMAP) for integrated analysis of SC and IN group cells yields 16 distinct cell clusters. Cluster numbers are assigned based on largest population (cluster 1) to smallest (cluster 16).

Integrated analysis of tetramer-positive cells from the IN and SC groups yielded 16 distinct cell clusters numbered in order of decreasing size (Fig. 1c, Supplemental Table 1). With exception of dividing populations described below, clusters contained cells across all stages of the cell cycle and were not affected by regression of cell cycle genes (Supplemental Fig. 1). All cell clusters expressed *Cd3d* (CD3), *Cd4* (CD4), and *Trac* (TCRα constant chain), consistent with the gating strategy to select for CD4^+^ T cells (Fig. 2a). Most clusters also bore markers of tissue residence (T_RM_) such as *Cd69* (CD69), the galectins *Lgals1* and *Lgals3*, and *Vim* (vimentin) (Fig. 2b, Supplemental Fig. 2) (Szabo *et al*., 2019). Two populations (clusters 9, 16) specifically expressed *Ccr7* and *Sell*, markers of resting naïve or T_CM_ T cells that can be associated with lymphocyte transit to the site of infection (Fig. 2c) (Debes et al., 2005; Szabo *et al*., 2019). These findings indicate the expected presence of CD4^+^ memory T cells specific to the vaccine antigen with both tissue-resident and migratory phenotypes.

**Figure 2.**
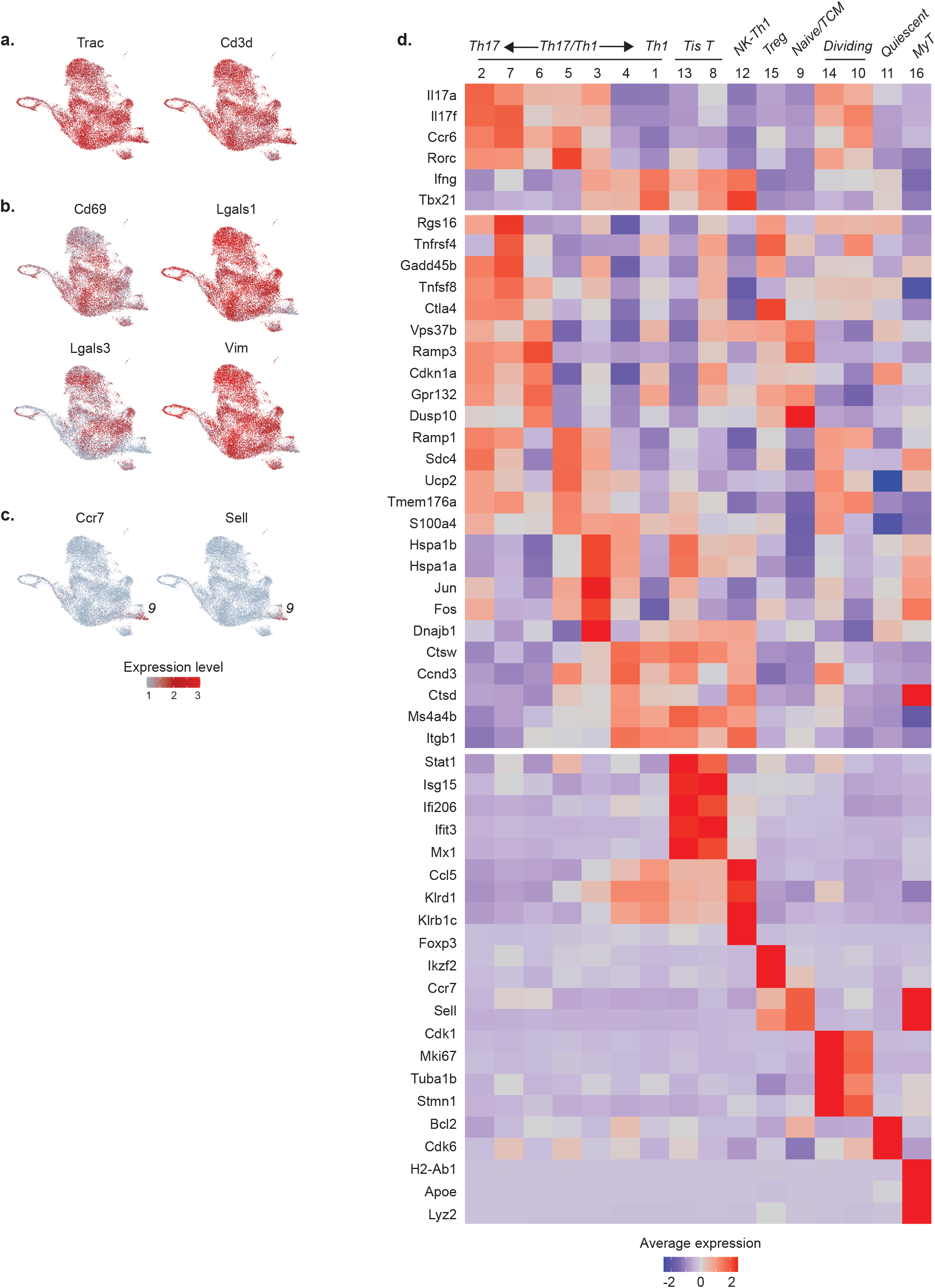
Identities of scRNAseq clusters. **(a)** Expression of CD4^+^ T cell genes (*Trac, Cd3d*) is consistent with gating strategy at the protein level across all 16 clusters. **(b)** UMAPs for T_RM_ markers *Cd69, Lgasl1, Lgals2*, and *Vim* show nonspecific expression patterns across most clusters. **(c)** UMAPs for resting (naïve/T_CM_) markers *Ccr7* and *Sell* show expression localizing to clusters 9 and 16. **(d)** Heatmap showing average expression and percent of cells expressing key genes to assign cluster identities. Marker genes include known lymphocyte marker genes and top differentially expressed genes identified as cluster markers by the scRNAseq package Seurat. Abbreviations: Tis T = Type I interferon signature T cells; NK-Th1 = NK-like Th1 cells; MyT = myeloid-like T cells.

### Cluster identities

Specific cluster identities were further interrogated by a combination of known marker genes and cluster markers identified by scRNAseq differential expression analysis (Fig. 2d). The largest populations bore conventional Th1 and Th17 cell signatures: high expression of *Ifng* (IFNγ) and Th1 transcription factor *Tbx21* (T-bet) (cluster 1); and high expression of *Il17a* (IL-17A), *Il17f* (IL-17F), *Ccr6*, and Th17 transcription factor *Rorc* (RORγt) (cluster 2). Interestingly, these clusters adjoined a spectrum of 5 additional populations also expressing Th1 genes, Th17 genes, or both (clusters 3-7). In addition to their cytokine phenotype, these clusters were distinguished by expression of genes less familiar to the classic Th framework, including: *Ctsw* and *Ctsd* (cathepsins W and D; cluster 4); *Vps37b* (vacuolar protein sorting 37B) and *Ramp3* (receptor activity modifying protein 3) (cluster 6) (Miragaia et al., 2019); co-stimulatory signaling genes *Tnfrsf4* (OX40/CD134) and *Tnfrsf9* (4-1BBL/CD137) (cluster 7); *Ramp1* (receptor activity modifying protein 1; cluster 5); and activator protein subunit genes *Jun* and *Fos* (cluster 3).

An unexpected and remarkable finding is that two populations (clusters 8, 13) expressed high levels of type I interferon response genes (*Stat1, Isg15, Ifi206, Ifit3, Mx1*). Although the type I interferon response is classically understood as an antiviral program, this type I interferon signature (Tis) has been described elsewhere outside of an antiviral immune context in CD4^+^ T helper cells, Tregs, and thymic invariant natural killer T (iNKT) cells (Andreatta *et al*., 2021; Arazi *et al*., 2019; Gowthaman *et al*., 2019; Harsha Krovi *et al*., 2020; Kiner *et al*., 2021; Seumois *et al*., 2020; Singhania *et al*., 2019; Szabo *et al*., 2019; Tibbitt *et al*., 2019; Zemmour et al., 2020). Here, we adopt the term Tis T cells to describe these distinct populations. We also observed a population (cluster 12) notable for very high expression of chemokine *Ccl5* (CCL5) and multiple NK-cell markers including *Nkg7* (natural killer granule protein 7), *Klrd1* (CD94), and *Klrb1c* (CD161). Since this cluster also expressed Th1 genes (*Ifng, Tbx21*) at a level comparable to a conventional Th1 phenotype (cluster 1), we provisionally termed this cluster NK-like Th1 cells.

The remaining populations included Tregs (*Foxp3, Ikzf2*; cluster 15), two populations of dividing cells (*Cdk1, Mki67, Tuba1b, Stmn1*; clusters 14, 10), and naïve/T_CM_ cells (*Ccr7, Sell*; cluster 9) (Szabo *et al*., 2019). We also saw a population of transcriptionally less active cells (cluster 11) that expressed markers for prolonged survival (*Bcl2, Cdk6*), suggesting quiescent cells distinct from the resting naïve/T_CM_ population expressing *Ccr7* and *Sell* (Supplemental Fig. 2) (Cheng et al., 2004). Lastly, we observed a very small population of cells bearing myeloid markers (*H2-Ab1, Apoe, Lyz2*; cluster 16). Since this smallest group of cells still expressed CD4^+^ T cell markers and did not exhibit increased reads suggestive of myeloid cell-T cell doublets (Supplemental Fig. 2), we tentatively labeled them myeloid-like T (MyT) cells, adopting the term for a population of αβ T cells that acquire myeloid markers peripherally and have been validated elsewhere with flow cytometry and RNAseq (Kiner *et al*., 2021). Our ability to validate this novel cell type, however, was limited by the small number of cells for analysis (N=44, 0.2% all cells).

### Differential abundance and gene expression between cells from IN and SC vaccinated mice

The relative abundance of many clusters differed between the IN and SC groups (Fig. 3a,b; Supplemental Table 1, Supplemental Fig. 1). IN populations with increased relative abundance included *Il17a*-producing clusters (clusters 2, 6, 7), naïve/T_CM_ cells (cluster 9), one population of dividing cells (cluster 14), and Tregs (cluster 15). By contrast, SC populations with increased relative abundance included *Ifng*-expressing clusters (clusters 1, 4), Tis T cells (clusters 8, 13), and NK-like Th1 cells (cluster 12). Unsurprisingly, the shift in relative abundance was associated with differential gene expression across all antigen-specific cells in the IN group compared to SC group (Fig. 3c; Supplemental Table 2). Average *Il17a* expression was higher for the IN group, consistent with a Th17-skewed response to pathogen challenge seen previously with this route of vaccination, as was expression of other intercellular signaling genes including *Ccr6* (CCR6) and *Cxcr4* (CXCR4) (Dobson *et al*., 2020). By contrast, the SC group showed higher average expression of *Ifng*, macrophage- and granulocyte/macrophage-stimulating genes *Csf1* (M-CSF) and *Csf2* (GM-CSF), the chemokine *Ccl5* (CCL5, aka RANTES), and chemokine receptor *Cxcr6* (CXCR6). This skewed Th17 response in the unprotected IN group was unexpected, since Th17 response is generally believed to promote protection against fungi at mucosal surfaces (Huppler et al., 2012).

**Figure 3.**
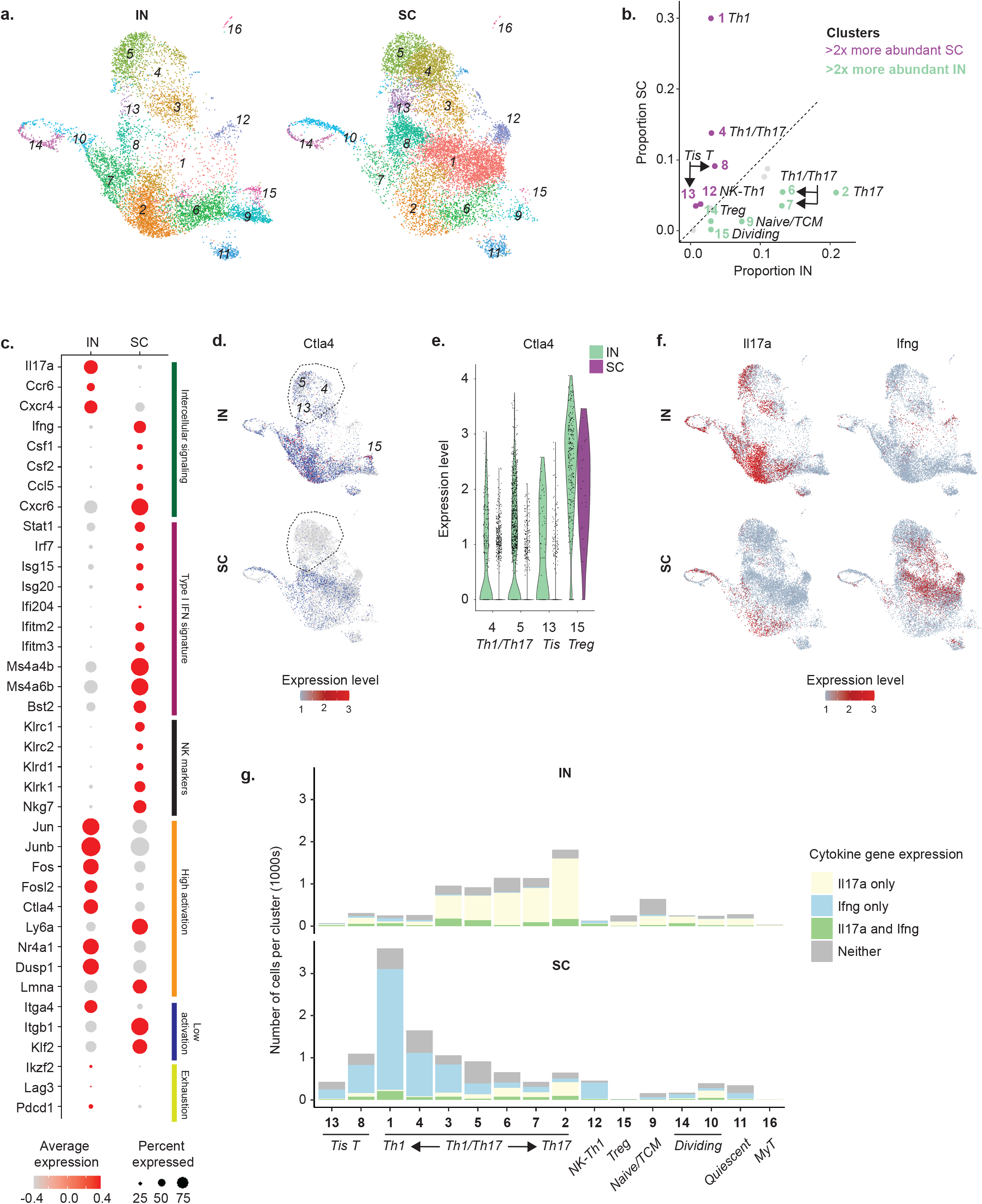
Differences in relative cluster abundance and gene expression between IN and SC vaccinated mice. **(a)** UMAP of 16 cell clusters separated by IN and SC sample origin, showing shifts in relative cluster abundance between these two groups. **(b)** Scatterplot of relative proportions of each cluster within all IN group cells (x-axis) and SC group cells (y-axis). Clusters falling above dotted line have higher relative abundance for SC group; clusters below dotted line have higher relative abundance for IN group. Those with >2x increased relative abundance in IN or SC samples are green and purple, respectively; cluster 16 made up <1% of all cells and was not color coded. **(c)** Dot plot of key genes with differential expression between IN and SC samples, grouped by functional similarity (colored annotation to the right of panel); all with exception of *Lag3* and *Pdcd1* are significantly different (*p* < 0.05). High and low activation gene lists are adapted from scRNAseq profiling of CD4^+^ T cells elsewhere (see Kiner et al., *e*.*g*. Fig 1d.). **(d)** UMAP for activation-induced immune chekpoint gene *Ctla4*. Circled clusters 4, 5 and 13 are shown in detail with the violin plot for *Ctla4* expression in **(e)**, where these clusters show decreased expression SC compared to IN. This difference contrasts with stable *Ctla4* expression in Tregs in both IN and SC groups. **(f)** UMAP for *Il17a* and *Ifng*, separated by IN and SC groups. **(g)** Number of cells within each cluster expressing *Il17a* only, *Ifng* only, both *Il17a* and *Ifng*, or neither cytokine. Note the switch in dominant cytokine from *Il17a* to *Ifng* within certain clusters (e.g. cluster 3) depending on vaccine exposure route.

Other salient differences included increased expression in the SC group of type I interferon response genes and NK cell markers, as would be expected with increased abundance of Tis T and NK-like Th1 cells. In the IN group, we also observed increased expression of some activation-related genes such as Jun and Fos family genes (*Jun, Junb, Fos, Fosl2*) and activation-induced immune checkpoint gene *Ctla4* (CTLA-4) (Kiner *et al*., 2021). Since high activation genes did not universally segregate to the IN group, nor low activation genes to the SC group, this data suggests complexity beyond the hypothesis that one vaccine route might prime a more activated CD4^+^ T cell phenotype than the other route during pathogen challenge. Nevertheless, our data showed a trend towards increased expression of some exhaustion markers such as *Ikzf2* (Helios), *Lag3* (Lymphocyte activation gene 3), and *Pdcd1* (Programmed cell death 1 [Pd1]) (Fig. 3c). While this difference did not correspond to marked differences between the IN and SC groups in the relative abundance of proliferating cells (Fig. 3b), we did observe increased expression of activation-induced immune checkpoint gene *Ctla4* in the IN group, which was not exclusively explained by the increased abundance of Tregs, but rather appeared to reflect specific downregulation of this critical checkpoint gene in non-Treg clusters in the SC group (Fig. 3d, e). This data may suggest greater activation-induced exhaustion in the IN group, or perhaps escape from this negative feedback mechanism in the protective vaccine-induced immune response.

### Analysis of Th1 and Th17 phenotypes

We next analyzed characteristic Th1 and Th17 cytokine gene expression at the single cell level. Interestingly, the Th17 cytokine phenotype in the IN group reflected not only the relative expansion of populations characterized by *Il17a* expression regardless of vaccination route (e.g. cluster 2), but also from a increased *Il17a* expression within populations that would otherwise express *Ifng* in mice vaccinated SC (e.g. cluster 3) (Fig. 3f). Similarly, the Th1 cytokine phenotype in the SC group appeared associated with increased abundance of populations restricted to *Ifng* expression (e.g. clusters 1, 4) as well as a switch from *Il17a* to *Ifng* dominance in other Th1/Th17 populations. The distribution of Th1 and Th17 transcription factors *Tbx21* (Tbet) and *Rorc* (RORγt) mirrored the patterns seen in downstream cytokine expression (Supplemental Fig. 2). This fluidity of the dominant cytokine phenotypes in Th1/Th17 cells complicates assignments of strict Th archetypes and may align with evolving notions of Th cell cytokine plasticity, for instance in Th17 cells described elsewhere (Zhu and Paul, 2010). Alternatively, this result may indicate dual cytokine production among cells from each vaccine group. There were stable small fractions of cells producing both cytokines simultaneously (Fig. 3f), though the limited sequencing depth for each cell in scRNAseq makes it challenging to differentiate whether these subpopulations are a true minority or simply undersampled.

### TCR gene usage

We explored whether cytokine phenotypes reflected the presence of a few expanded, highly active T cell clones, or a broader diversity of T cells responding to pathogen challenge. Using TRUST4, an algorithm that infers TCR clonotypes using focused reconstruction of variable TCR gene regions (Song et al., 2021), we recovered sufficient TCR sequence data from all 16 clusters to assign clonotypes by α, β, or combined αβ TCR sequences to 2,421 and 2,619 T cells in the IN and SC samples, respectively (Supplemental Fig. 3). Especially for α and αβ chains, we observed that clonotypes tended to be skewed in distribution between the IN and SC groups, with individual clonotypes occurring predominantly in either one or the other group (Fig. 4a). The IN group showed dominance of relatively few clonotypes, while the SC group exhibited more even representation of clonotypes comprising at least 2% of either sample (Fig. 4b, Supplemental Fig. 3). In both groups, the most frequent clonotypes were most abundant in the largest clusters expressing high levels of Th1 and Th17 cytokines (*e*.*g*., clusters 1-7; Fig. 4c, Supplemental Fig. 3). Thus, the size of these clusters appeared to reflect expanded, active T cell clones, with many TCR sequences unique to either the IN or SC group.

**Figure 4.**
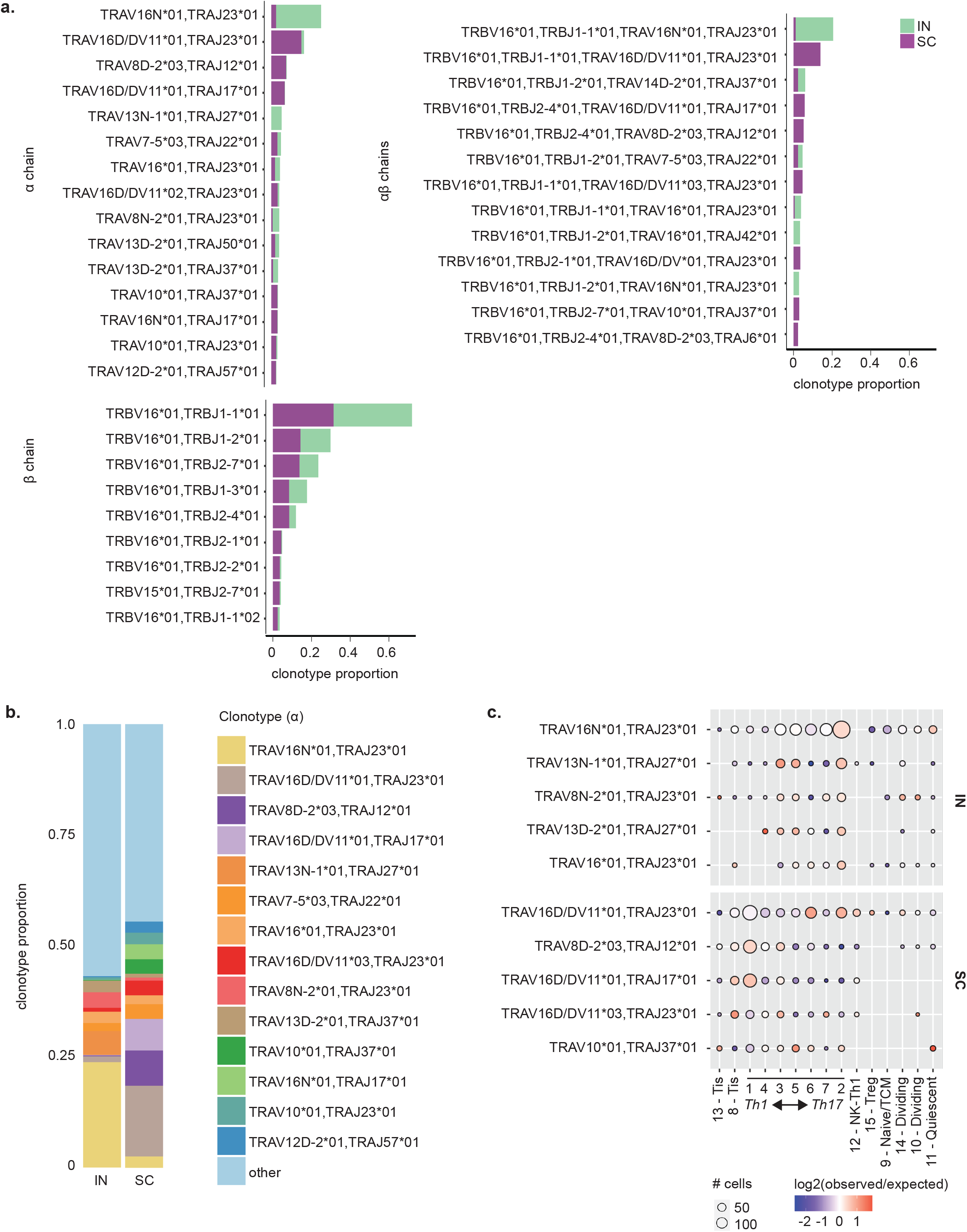
TCR clonotype diversity in IN and SC vaccinated mice. **(a)** Relative abundance of TCR α, β, and αβ chain clonotypes identified by TRUST4 in IN and SC samples. **(b)** TCR α-chain clonotypes vary in relative abundance between IN and SC samples. Only clonotypes with >2% abundance in either sample are shown. **(c)** Distribution of the top 5 TCR α-chain clonotypes across each cluster. The size of each circle depicts the number of cells expressing each clonotype per cluster. The color represents how relatively enriched (red) or depleted (blue) that cluster is for a given clonotype. White color indicates that the frequency of the clonotype is the same as the frequency across the entire sample.

### Special populations in nonprotective immune response: Tregs

We noted the higher relative abundance of *Foxp3*-expressing Tregs in the nonprotective IN vaccine-induced immune response (Fig. 5a), consistent with prior experimental data in this model (Dobson *et al*., 2020). Since mucosal antigen exposure can induce systemic immune tolerance, we wondered whether these Tregs might be impairing pathogen clearance by actively suppressing antifungal immunity (Rezende and Weiner, 2017). Tregs in the IN group, however, did not express high levels of tolerogenic cytokine genes such as *Tgfb1* (transforming growth factor β), *Il10* (interleukin-10), or *Il4* (interleukin-4) (Fig. 5b). Nor did we observe increased expression of markers of T cell anergy (e.g. *Rnf128* [GRAIL]) to suggest other mechanisms of tolerance in cells of the IN group (Supplemental Fig. 4). Indeed, other observed features of the response to pathogen challenge in the IN group, including prominent *Il17a* expression, appeared more consistent with pro-inflammatory response to a lethal pathogen than with microbial tolerance.

**Figure 5.**
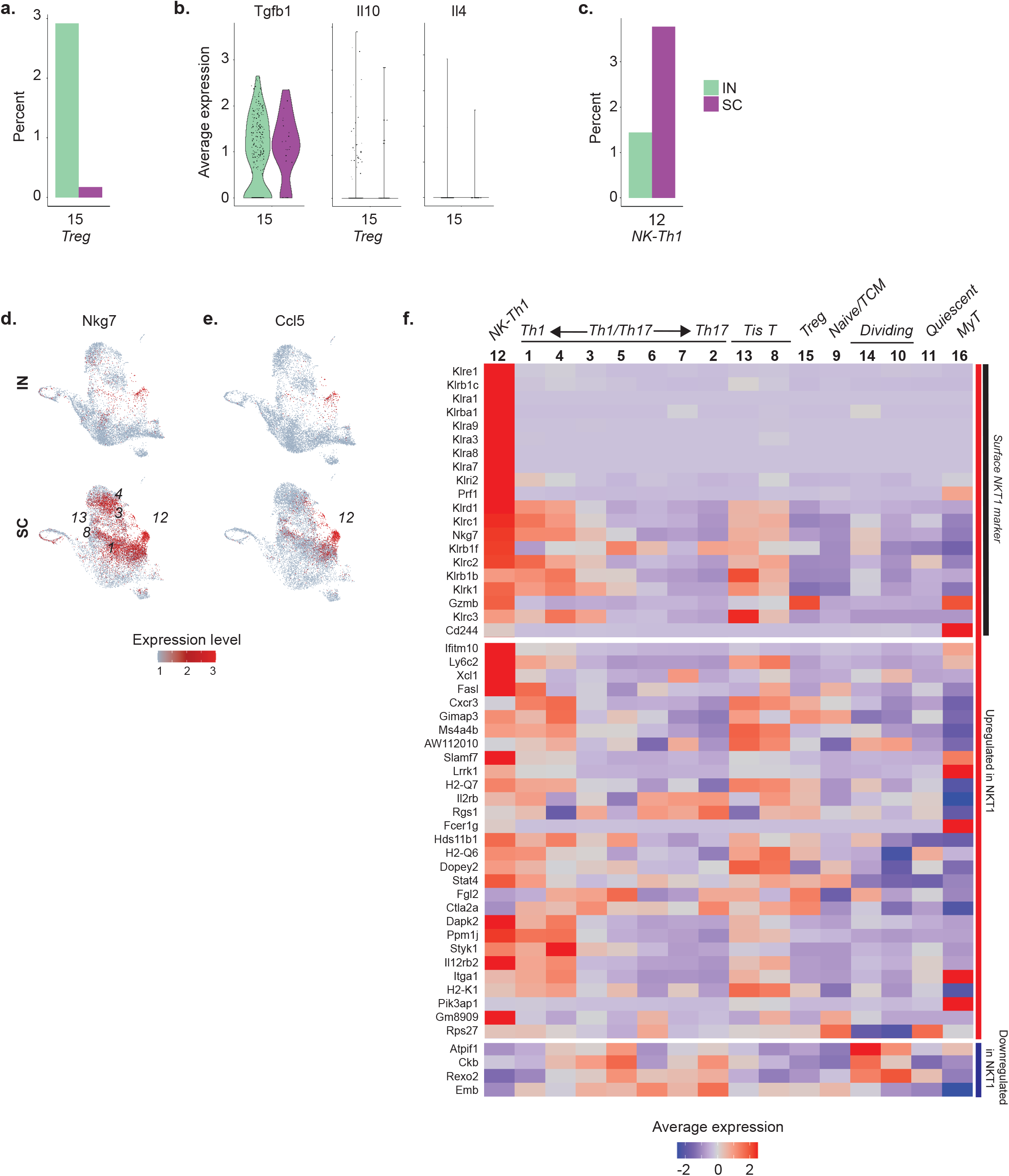
Tregs and NK-like Th1 cells in IN and SC vaccinated mice. **(a)** Relative abundance of Tregs in IN and SC samples, expressed as percent of total cells in each sample. **(b)** Violin plot showing average expression of tolerogenic Treg cytokines *Tgfb1* (TGFβ), *Il10* (IL-10), and *Il4* (IL-4), IN compared to SC. **(c)** Relative abundance of cluster 12 cells (NK-like Th1) in IN and SC samples, expressed as percent of total cells in each sample. **(d)** UMAP for *Nkg7*, an NK marker highly expressed in multiple SC-enriched populations including cluster 12. **(e)** UMAP for chemokine *Ccl5*, showing focal expression by cluster 12 cells. **(f)** Heatmap of average cluster expression of NKT1 marker genes identified in thymic Cd1d tetramer-positive cells (see: Engel et al., Figure 5).(Engel *et al*., 2016) Markers are primarily upregulated in NKT1 cells but include a subset of 4 genes downregulated in NKT1 (*Atpif1, Ckb, Rexo2, Emb*).

### Special populations in protective immune response: NK-like Th1 cells

We explored whether populations specific to the SC group might explain the distinctive efficacy of this vaccine immune response. One such population was the NK-like Th1 cells (cluster 12), for which a gene set enrichment analysis of its expression profile relative to cells from all other clusters resulted in the NK cell type as the most significantly enriched mouse cell type signature (adjusted p-value < 3 x10^−9^). This cluster was distinct for high levels of NK markers such as *Nkg7* (natural killer granule protein 7), *Klrb1c* (CD161), and *Klrd1* (CD94) (Fig. 5c). Among these markers, *Nkg7* raised particular interest due to an emerging role for inducible *Nkg7* in CD4^+^ T cells, where it appears to be associated with IFNγ expression and promote parasite control in a model of *Leishmania donovani* infection (Ng et al., 2020). Indeed, we observed increased *Nkg7* expression in those populations enriched in the SC group’s *Ifng-*polarized immune response, including in the highest *Ifng*-expressing populations (clusters 1, 4, 3), Tis T cells (clusters 13, 8), and NK-like Th1 cells (Fig. 5d). These NK-like Th1 cells were also notable for specific expression of the gene *Ccl5* (CCL5 or RANTES), a pleiotropic chemokine that attracts effector and memory cells to the site of infection and is unique among CC-type chemokines for its role in the later stages of response to infection (Fig. 5e) (Ortiz et al., 1996). Notably, while NK markers such as granzyme (*Gzmb*) and perforin (*Prf1*) are associated with cytotoxic function, our NK-like Th1 cells did not express markers of cytotoxic CD4^+^ T cells, a recently described population that appears capable of inducing apoptosis of target cells in an MHC class II-restricted manner (Supplemental Fig. 4) (Takeuchi and Saito, 2017).

Based on NK markers, we considered whether these cells might be NKT cells, an innate-like T cell population that expresses αβTCR and combines NK cell reactivity with some of the antigen-specificity of T cells (Godfrey et al., 2004). Indeed, cluster 12 cells expressed many of the genes up- and down-regulated in Th1-like NKT cells (NKT1) by scRNAseq profiling (Fig. 5f) (Engel et al., 2016). Importantly, however, these cells lacked expression of the gene for PZLF (*Zbtb16*), a transcription factor marker for most innate and innate-like T cell populations including NKT cells (Supplemental Fig. 4) (Mao et al., 2017). Moreover, the TCR of NKT cells classically binds Cd1d, an MHC class I-type receptor that presents lipid antigen, and would not be expected to bind the MHC class II tetramer and peptide antigen used to sort our Bl-Eng2-specific T cells. While some have reported NKT cells in CD1d-deficient mice—including CCL5 producers as seen here—others insist on CD1d-restriction as an essential feature for the term NKT to remain meaningful (Eberl et al., 1999; Farr et al., 2014; Giroux and Denis, 2005; Godfrey *et al*., 2004). We opted for the term NK-like Th1 cells, emphasizing core Th1 features with additional NK marker expression. In either case, the appearance of this NK-like Th1 phenotype and the accompanying chemokine activity were salient, previously undescribed features of vaccine-induced protective immunity to fungi.

### Special populations in the protective immune response: Tis T cells

Tis (type I interferon signature) T cells (clusters 8, 13) were another cell phenotype associated with the protective immune response. Especially in the SC group, these cells comprised a meaningful portion of our tetramer positive T cells, representing 12.6% and 4.2% of all cells in the SC and IN vaccine groups, respectively (Fig. 6a). These populations showed a transcriptional signature dominated by several type I interferon-responsive genes (*Ifitm3, Ifi204, Isg15, Isg20, Mx1, Rsad2, Oas3*, etc.) (Fig. 6b). This signature included genes upstream in type I interferon signal transduction, such as *Stat1* (Stat1) and *Stat2* (Stat2), and those associated with distal interferon response functions such as global suppression of translation (*e*.*g. Eif2ak2* [EIF2α kinase 2]), processing of cytosolic DNA and RNA (e.g. *Ddx58* [RIG-I], *Zbp1* [Z-DNA binding protein 1], and *Samhd1* [SAM and HD domain 1]), and protection from viral infections including influenza and SARS-CoV-2 (*Ifitm3* [interferon-induced transmembrane protein 3]) (Fig. 6c, Supplemental Table 3) (Everitt et al., 2012; Prelli Bozzo et al., 2021). Other Tis T cell genes coding for transmembrane proteins (*Rtp4, Bst2* [tetherin/CD317]) and nuclear body proteins (*Pml, Sp100*) were noteworthy as potential cell surface or microstructural markers.

**Figure 6.**
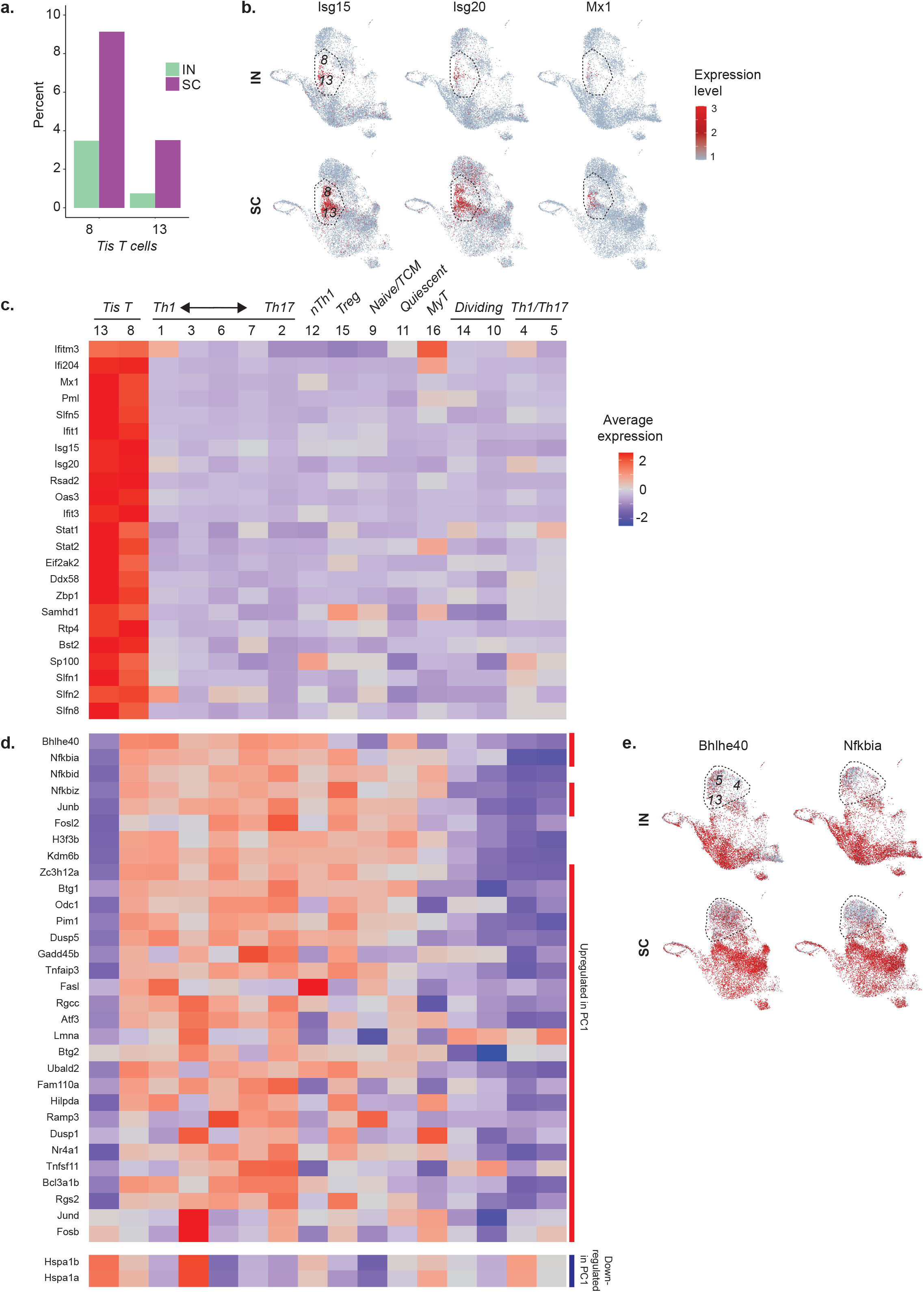
Tis T cell phenotypes in IN and SC vaccinated mice. **(a)** Relative abundance of Tis T cells in IN and SC samples, expressed as percent of total cells in each sample. **(b)** UMAP for expression of type I interferon response genes, localizing Tis T cells to clusters 8 and 13. **(c)** Heatmap showing average expression and percent of cells expressing Tis T cell markers noted in the literature (*e*.*g. Isg15, Isg20, Mx1, Rsad2, Oas3, Ifit1, Ifit3*) and identified as markers for both clusters by scRNAseq differential expression analysis. **(d)** Heatmap for genes differing between the two clusters of Tis T cells. Red and blue side bars represent those genes that are upregulated or downregulated in the first principal component (PC1) of CD4^+^ T cells harvested after infection with enteric pathogens (*Salmonella typhimurium, Citrobacter rodentium, Heligmosomoides polygyrus* and *Nippostrongylus brasilensis*; see Kiner, Supplemental figure 2) (Kiner, 2019). **(e)** UMAP showing similar patterns in activation gene expression, represented by *Bhlhe40* and *Nfkbia*, between cluster 13 Tis T cells and nearby clusters 4 and 5, both IN and SC.

### Tis T cell heterogeneity

Tis T cells separated into 2 clusters that shared a common strong type I interferon signature (Fig. 6c). However, one Tis T cell population (cluster 8) distinctly expressed more *Bhlhe40* (basic helix-loop-helix family member e40), a key transcription factor that characterizes a highly pro-inflammatory phenotype in CD4^+^ memory T cells (Fig. 6d, Supplemental Table 4) (Emming et al., 2020). This same Tis T cell cluster also expressed higher levels of NF-κB inhibitors *Nfkbia* (IκBα), *Nfkbid* (IκBNS), and *Nfkbiz* (IκBζ) (Emming *et al*., 2020), activator protein 1 (AP-1) subunit genes *Junb* (Junb) and *Fosl2* (Fra2), histone and histone modulating genes (e.g. *H3f3b* [H3.3 histone B], *Kdm6b* [lysine demethylase 6B]), and *Zc3h12a* (MCPIP1 or regnase-1) (Garg et al., 2015; Matsushita et al., 2009). Remarkably, many of these same markers have appeared recently in another scRNAseq analysis of CD4^+^ T cell heterogeneity, in which transcriptional diversity was driven primarily by activation state rather than by conventional Th archetype as might have been expected (Kiner, 2019; Kiner *et al*., 2021). In that analysis, activation-related genes (*e*.*g. Bhlhe40, Jund, Dusp1, Btg1, Odc1, Vps37b*) comprised the first principal component (PC1) driving transcriptional diversity in CD4^+^ T cells following a variety of enteric infections (Kiner, 2019). In querying our data for these PC1 genes, we saw that differing expression not only separated our Tis T cell populations, but also organized non-Tis T cells into two rough superclusters, in which Th1/Th17 clusters 4 and 5 grouped with cluster 13 Tis T cells apart from surrounding Tis and non-Tis T cells (*e*.*g. Bhlhe40, Nfkbia*; Fig. 6e). This finding emphasized the importance of activation state as an organizing principle for CD4^+^ T cell heterogeneity, including within Tis T cells.

We considered whether the separation of Tis T cells into two populations (clusters 8 and 13) might reflect a sequence of cell differentiation, in which one phenotype might be a precursor to the other. These transitional patterns can be explored in scRNAseq with RNA velocity analysis. The ratio of unspliced and spliced reads mapping to a given gene is compared to expected steady state kinetics to make a prediction about an increase in transcription rate (with the resulting increase in unspliced mRNA) or vice versa (La Manno et al., 2018). In our Tis T cells, however, we did not observe any genes characterizing the expression profile of one cluster among the genes with most significantly different velocity in the other (Supplemental Table 5). This finding suggests that one population is not a precursor of the other population.

### Validation of Tis T cell phenotype

We validated the presence of the Tis T cell phenotype by quantitative RT-PCR in vaccinated mice after *Blastomyces* challenge. Consistent with scRNAseq data, we detected a transcriptional signal for multiple Tis T cell marker genes (*Ifi204, Mx1, Pml, Slfn5, Ifit1, Ifitm3*) present among tetramer^+^ antigen-specific CD44^+^ cells, but not among control (CD44^-^) cells, in both the lung and spleen after pulmonary pathogen challenge (Fig. 7a, Supplemental Fig. 5). This difference in expression was distinct from the upregulation of interferon response genes in both antigen-specific and control cells following exposure to soluble type I interferon (IFNα), though the relative enrichment of several Tis T cell transcripts in antigen-specific cells relative to control cells was still detectable in this experiment. As expected, relative expression of Tis T cell markers *Ifi204* and *Ifitm3* was increased in tetramer-positive T cells from SC vaccinated animals compared to those from the IN group (Fig. 7b), conforming to results from our scRNAseq analysis (Fig. 3b).

**Figure 7.**
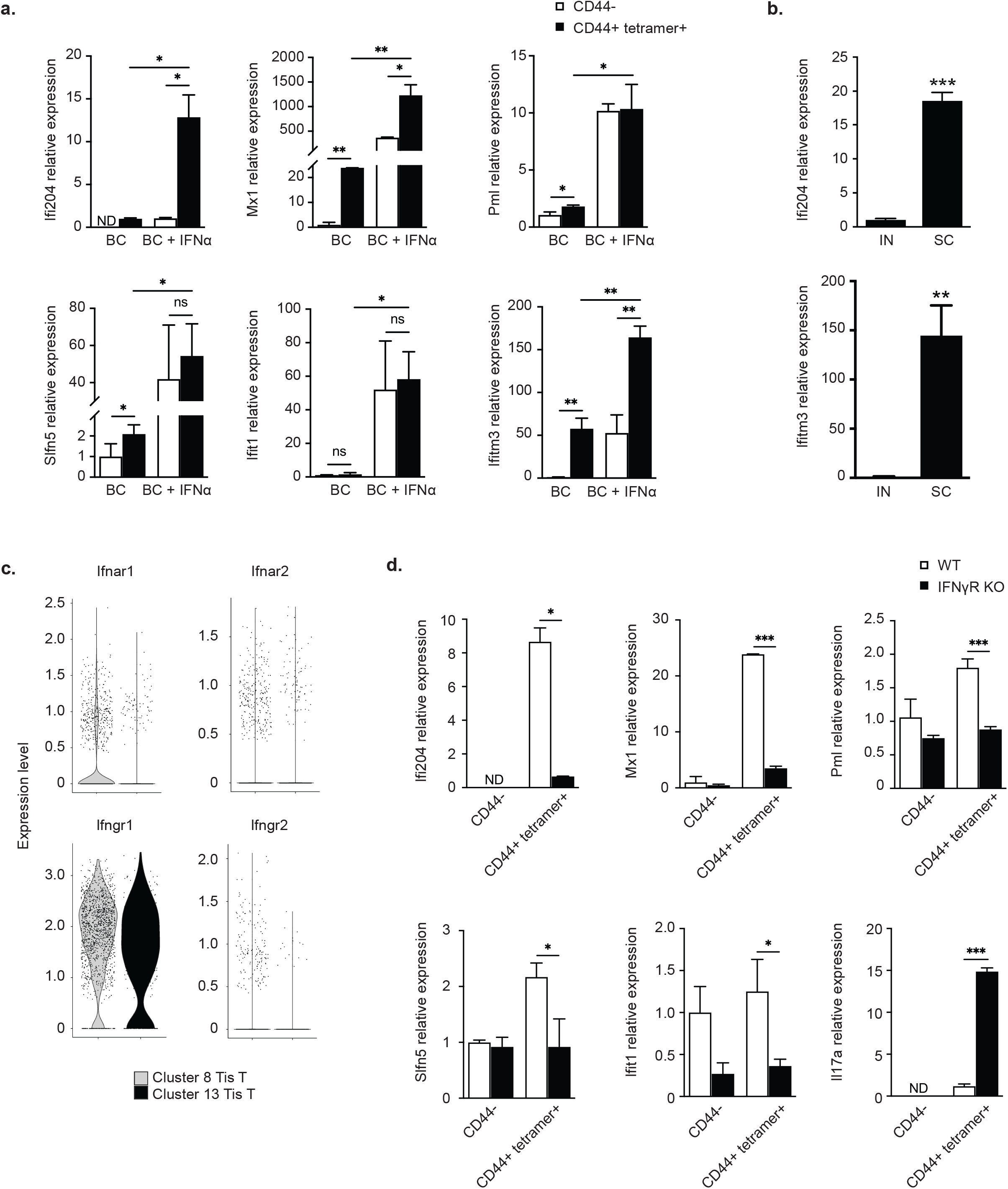
RT-PCR Tis T cell signature. **(a)** Quantitative reverse transcriptase PCR (RT-qPCR) detects increased expression of Tis T cell marker genes (*Ifi204, Mx1, Pml, Slfn5, Ifit1, Ifitm3*) in tetramer-positive, CD44-positive (black) cells compared to control CD44-negative (white) cells from the lungs of *Blastomyces*-challenged, subcutaneously vaccinated mice. The addition of IFNα increased expression of these genes in both tetramer-positive and control cells. **(b)** Tis T cell signature (*Ifi204, Ifitm3*) was increased in SC vaccinated animals compared to IN vaccinated animals by RT-qPCR. **(c)** Violin plot depicting expression of interferon receptor genes in Tis T cells (clusters 8, 13). While these clusters expressed the IFNγR α-chain, the corresponding β-chain was not detected. **(d)** Tis T cell signature is diminished in lungs of *Blastomyces*-challenged, subcutaneously vaccinated mice lacking IFNγR (IFNγR-KO) as compared to wild type mice. *, p<0.05; **p<0.01; ***, p<0.001. Analysis by two-way ANOVA.

### IFNγR-dependence of Tis T cell signal

We sought to understand upstream signaling for the Tis T cell signature. Remarkably, despite an increased and highly specific type I interferon gene signature, our Tis T cell populations did not express the type I interferon receptor genes *Ifnar1* (IFNAR1) and *Ifnar2* (IFNAR2) (Fig. 7c) (Tibbitt *et al*., 2019). Other investigators studying *T. gondii*-infected mice have described a strong type I IFN transcription module dependent on the presence of IFNγR, another type II cytokine receptor (Singhania *et al*., 2019). We hypothesized that this transcription module might reflect the presence of Tis T cells and tested whether our Tis T cell signature may similarly depend on IFNγR. To explore this idea, we vaccinated IFNγR^-/-^ (IFNγR KO) mice SC. We found that, after pulmonary challenge, the presence of signature transcripts for Tis T cells was diminished in tetramer^+^ antigen-specific T cells, but not in CD44^-^ control cells (Fig. 7d). Thus, the emergence of Tis T cells requires IFNgR signaling. In contrast to the Tis T cell transcripts, the relative expression of *Il17a* was increased in IFNγR KO mice, suggesting a compensatory effect in the dynamic balance between Th1 and Th17 cytokine environments. Despite this potential compensation, IFNγR KO did not acquire resistance after SC vaccination. After pulmonary challenge, the lungs of these mice were grossly abnormal. They were more swollen than the lungs of corresponding wild-type mice and with nodules and micro-abscesses stippling the pleural surface. This increased inflammation was accompanied by increased percentages of CD4^+^ T cells and tetramer-positive cells in the lungs of the IFNγR KO mice compared to wild type mice (Supplemental Fig. 5d)

Intriguingly, while scRNAseq data did show that Tis T cells express IFNγR α-chain gene (*Ifngr1*), these cells did not express the IFNγR β-chain gene (*Ifngr2*) presumed to confer IFNγ responsiveness to T cells (Fig. 7e) (Bach et al., 1997; Bach et al., 1995). Moreover, soluble IFNγ did not elicit any upregulation of the Tis T cell signature genes in tetramer positive cells in our RT-PCR experiments (Supplemental Fig. 5). Together, this data suggests the dependence of the Tis T cell signature on IFNγR, apparently by a mechanism distinct from classic IFNγ-IFNγR receptor signaling.

## DISCUSSION

Antigen-specific T cells are essential for vaccine-induced immunological memory and effective pathogen control. Our work demonstrates key differences in the phenotypic profiles of antigen-specific CD4^+^ T cells present at the site of pathogen challenge in a protective and nonprotective vaccine model. Our study yields some distinctly surprising results in addition to expected findings. We observe unique cell populations, including two with high expression of type I interferon signature genes and one highly expressing *Ccl5*, that are associated specifically with protective vaccine-induced immunity. We also redemonstrate essential differences in Th1- and Th17-skewing of protective and nonprotective vaccine-induced responses, reflecting the activity of conventional appearing, clonally dominant Th1 and Th17 cells together with a spectrum of more phenotypically heterogeneous *Ifng*- and *Il17a*-expressing populations.

We uncovered a Tis T cell phenotype enriched among antigen-specific cells that confer vaccine-induced immunity. We did not expect to see type I interferon signaling in the context of vaccine-primed responses to *B. dermatitidis*, a fungal pathogen traditionally understood to elicit Th1- and Th17-related cytokines such as IFN-γ, TNF-a, IL-17, and IL-6 (Merkhofer et al., 2019; Speakman et al., 2020). Others have recently observed Tis T cells in settings outside of viral infection (Andreatta *et al*., 2021; Arazi *et al*., 2019; Gowthaman *et al*., 2019; Harsha Krovi *et al*., 2020; Kiner *et al*., 2021; Seumois *et al*., 2020; Singhania *et al*., 2019; Szabo *et al*., 2019; Tibbitt *et al*., 2019; Zemmour *et al*., 2020). However, our findings—validated by RT-qPCR—add new insight to this enigmatic, recently described population of T cells. In our model, Tis T cells were associated with a protective, vaccine-induced immune response. We also observed phenotypic heterogeneity underlying the Tis T cell signature that has not been previously described, including divergent expression of the pro-inflammatory transcription factor *Bhlhe40* and other activation-related genes that appear to be an organizing framework for CD4^+^ T cells across microbially diverse infectious challenges (Emming *et al*., 2020; Kiner *et al*., 2021). The loss of the Tis T cell signature in IFNγR KO mice is another remarkable feature that merits future study. We wonder whether the IFNγR α-chain expressed nonspecifically in Tis T cells could bind another unidentified cytokine receptor chain required for a novel type I IFN signature response, analogous to the combinatorial plasticity seen in other cytokine signaling mechanisms (Morris et al., 2018). Alternatively, the loss of IFNγR could indirectly mute the Tis T cell response, e.g. through decreased T cell activation in IFNγR KO mice overall. These hypotheses and other features of Tis T cell biology, such as the possible antagonism of Tis T cells by Th17 cells within the IN group, are exciting avenues for future functional studies.

Our findings highlight the difficulties of classifying populations in scRNAseq that share features with multiple conventionally defined cell types, which are often described by only a few markers. In our data, NK-like Th1 cells make up one such population that resists straightforward identification based on overlapping features with Th1, NK, and NKT cells. Nonetheless, this ambiguity will be important to pursue. This population is a distinct feature of a protective vaccine-induced immune response in our model and the sole source of *Ccl5* expression. CCL5 is known to be unique among CC chemokines as a late-appearing signal, expressed three to five days after T cell activation, with a role in attracting effector T cells and new memory T cells to the site of infection (Ortiz *et al*., 1996; Seo et al., 2020). The high expression of *Ccl5* three days after pathogen challenge in only the SC vaccine group might reflect a mechanism of improved pathogen clearance by early cytokine production and effector cell recruitment following protective vaccination. These cells also highly express *Nkg7*, a feature shared with other *Ifng*-expressing populations in the SC vaccinated group. The potential association between *Nkg7* expression and IFNγ activity aligns with new data linking *Nkg7* and CD4^+^ T cell activation and suggests a role for this molecule in regulating key effector cytokines from CD4^+^ T cells in a protective vaccine-induced immune response (Ng *et al*., 2020).

Lastly, our analysis confirms the core distinction between IFNγ- and IL-17-skewed responses in protective and nonprotective vaccine-induced immune responses, respectively (Dobson *et al*., 2020). Notably, the highest cytokine producing, clonally dominant Th1 and Th17 cells that express archetypical Th markers lie at the extremes of a spectrum of phenotypes that differ by expression of genes that are distinctly unfamiliar to a classic Th paradigm. For instance, among the 29 immune cell types of the Monaco dataset shared in the Human Protein Atlas, the cluster 7 marker *Rgs16* appears more specific to B cells than either Th1 or Th17 cells, and cluster 3 marker *Dnajb1* is similarly expressed in Th1, Th2, and Th17 cells (Monaco et al., 2019; The Human Protein Atlas, 2019). This reflects a tension between the widely accepted framework of conventional T helper cell archetypes (Th1, Th2, Th17) and the nuance that can be detected by newer, hypothesis-free approaches to immune cell profiling. More work remains to discern whether this heterogeneity is functionally meaningful—and if so, how it should be integrated into an organizing principle that remains useful for understanding immune cell ontogeny. Of note, these *Ifng*- and *Il17a*-expressing cell populations also appear to comprise largely non-overlapping TCR clonotypes, which might reflect either the stochastic effects of random V(D)J recombination prior to TCR selection by vaccination and pathogen challenge, some more active enrichment of specific TCR sequences within the protective immune response, or a combination of both.

Overall, our high-resolution single cell analysis of antigen-specific T cells in pathogen challenge provides insight into multiple dimensions of vaccine-related T cell biology. A general limitation of scRNAseq data is depth of sequencing, which comes at the cost of sequencing large numbers of cells (Zhang et al., 2020). More reads (*e*.*g*. greater depth) significantly reduces inaccuracy in estimating the true transcriptional state of a cell, but sequencing of more cells enables a broader view of the biological variability in the cell population. Consequently, we recognize that the absence of certain sequences does not exclude low level expression that was undetectable in our analysis. Other limitations inherent to study design include the lack of transcriptional data for antigen-nonspecific cells and of functional data for populations identified by scRNAseq. While our study uncovered correlations between novel populations of CD4 T cells and resistance, one should not assume causal relationships from the observed associations between cell phenotypes (*e*.*g*. Tis T cells, NK-like Th1 cells) and biological outcomes such as improved pathogen control following SC vaccination. Future studies are required to discern whether Tis T cells, CCL5, or NKG7 are required for protective vaccine-induced immunity or are simply markers of this response that is driven by other cellular events. Our analysis of TCR sequences is also limited by use of conventional 3’ library preparation, which provides less coverage of hypervariable regions clustered towards the 5’ end and may be reason for choosing 5’ chemistry for more robust TCR analyses and clonotype tracking in the future. Nonetheless, our work describes novel characteristics of vaccine-induced T cells in protective immunity, including populations that could serve as correlates of efficacy in vaccine design, and adds to the ongoing, exciting scientific pursuit of T cell diversity.

## METHODS

### Mice

*C57BL/6* mice from Jackson Laboratory were bred at our facility and cared for per guidelines from the University of Wisconsin Animal Care Committee, who approved all aspects of this work. Mice were 7-8 weeks old at the start of experiments. Mice were vaccinated intranasally (IN) or subcutaneously (SC) with 10 μg of Bl-Eng2 in glucan chitin particles (CGP) a total of three times, two weeks apart. Two weeks after the final vaccination, mice were challenged intratracheally with 2×10^4^ *Blastomyces dermatitidis* (*Bd*, ATCC strain 26199) and analyzed at day 3 post-infection.

### Flow cytometry

We harvested cells from a total of 24 mice: 10 vaccinated SC and 14 vaccinated IN. Cells were prepared from harvested lungs as described previously and pooled for each group.(Dobson *et al*., 2020) Briefly, lungs were harvested from challenged animals and dissociated in Miltenyi MACS tubes (Miltenyi Inc., Germany) and digested with collagenase (1 mg/mL) and DNase (1 µg/mL) for 25 min at 37 °C. Digested lungs were resuspended in 5 mL of 40% percoll, and 3 mL of 66% percoll was underlaid (GE healthcare 17–0891–01). Samples were spun for 20 min at 2000 rpm at room temperature. Lymphocytes were then harvested from the buffy coat layer and resuspended in complete RPMI (10% FBS, 1% penicillin and streptomycin). The cells were spun down (1500 rpm/5 minutes at room temperature) and stained with LIVE/DEAD™ Fixable Near-IR Dead Cell Stain Kit (Invitrogen) and Fc Block (BD) for 10 min at room temperature. Then the cells were stained with Bl-Eng2 tetramer (MHC class II tetramer-PE, NIH) for 1 hour at room temperature, and 30 minutes at 4°C with the following surface antibodies: CD8 PerCP-Cy5.5 (clone 53–6.7, Biolegend, cat#100734), CD44 BV650 (clone IM7, Biolegend, cat#103049), CD11b APC (clone M1/70, Biolegend, cat#101212), CD11c APC (clone N418, Biolegend, cat#117310), NK1.1 APC (clone PK136, Biolegend, cat#108710), B220 APC (clone RA3– 62B, Biolegend, cat#103212), CD4 BUV737 (clone RM4–5, BD, cat#565246), and CD90.2 BV421 (clone 30-H12, Biolegend, cat105341). All panels included a dump channel to decrease background in CD4+ T cells (Dump: CD11b, CD11c, NK1.1, and B220). The cells were sorted using the cell sorting flow cytometer FACSAria (BD). Following fluorescent labeling, cells from 10-15 animals from each vaccine (SC or IN) group were combined into one tube each for cell sorting. Tetramer^+^ cells were sorted into microcentrifuge tubes containing RPMI media on a FACs Aria using a 130 micron nozzle. The sorted cells (Live, Dump^-^CD90.2^+^CD4^+^CD44^+^Tetramer^+^) were collected directly into 1.5 ml microtubes and provided to the UW-Madison Biotechnology Center for 10x Genomics Single Cell RNA sequencing.

### Single-cell RNA-seq libraries

Sorted tetramer^+^ cells were counted on a Countless II cell counter with 0.4% trypan blue and concentrated to 300-400 cells/ml (total volume of 43.3 ml) and reverse transcribed. The libraries were generated with the 3’ kit version 3.1 chemistry (10x Genomics) and sequenced on the MiSeq system and the NovaSeq 6000.

### Single-cell RNA-seq data analysis

Single cell RNAseq data was initially processed by the UW Bioinformatics Resource Center. Experiment data was demultiplexed using the Cell Ranger Single Cell Software Suite, mkfastq command wrapped around Illumina’s bcl2fastq (v2.20.0.422). The MiSeq balancing run was quality controlled using calculations based on UMI-tools (Smith et al., 2017). Samples libraries were balanced for the number of estimated reads per cell and run on an Illumina NovaSeq system. Cell Ranger software version 3.1.0 was then used to perform demultiplexing, alignment, filtering, barcode counting, UMI counting, and gene expression estimation for each sample according to the 10x Genomics documentation (https://support.10xgenomics.com/single-cell-gene-expression/software/pipelines/latest/what-iscell-ranger). The reference for alignment was the curated 10x genomics reference for mouse (mm10-3.0.0). The gene expression estimates from each sample were then aggregated using Cellranger (cellranger aggr) to compare experimental groups with normalized sequencing-depth and expression data.

Single-cell expression data was then analyzed using Seurat 4.0 (Hao et al., 2021). Genes detected in fewer than 5 cells were filtered out of analysis. Doublets were removed from analysis, and cells with <2000 or >20,000 unique molecular identifiers (UMI) or <1000 or >3000 genes were excluded from analysis. Cells with elevated percentage of mitochondrial reads (>5%) were also excluded as a means to filter out dying cells. Ultimately 70.0% of cells IN and 80.1% of cells SC passed quality control filters. Data were normalized using the NormalizeData function, and IN and SC samples were integrated for downstream analysis using FindIntegrationAnchors and IntegrateData functions with methods described previously (Stuart et al., 2019). Clustering and visualization for the integrated dataset proceeded with a standard scRNAseq workflow including ScaleData, RunPCA, RunUMAP, FindNeighbors and FindClusters functions. FindClusters was run with resolution parameter 0.83 to achieve clusters that separated cell populations with previously established markers. Cluster markers were obtained with FindMarkers function (min.pct = 0.25). Dimension-reduced plots were generated with FeaturePlot function, splitting by original sample identity as needed for specific analyses. Heatmaps were produced with DoHeatmap function using cluster averages across both experiments calculated with AverageExpression function. Bar plots and scatterplots were generated using R package ggplot2. Gene set enrichment analysis was performed using the clusterProfiler R package using the CellMarker set of mouse cell type markers (Wu et al., 2021; Zhang et al., 2019). Specifically, the clusterProfiler function “GSEA” was run using a list of genes sorted by descending log2 fold change from the comparison of cells in one cluster vs. all others, as calculated by the Seurat FindMarkers function (logfc.threshold = 0). Scoring and prediction of cell cycle stage was performed using the Seurat CellCycleScoring function, with the lists of S and G2M genes provided by Seurat.

### TCR usage analysis

TCR sequence data was analyzed using TRUST4 v1.0.4.(Song *et al*., 2021) TRUST4 was run on the position sorted BAM file for each sample generated by CellRanger, along with the V/D/J/C gene reference files provided by TRUST4 (“GRCm38_bcrtcr.fa” and “mouse_IMGT+C.fa”), and the “--barcode CB” option to make clonotype calls for individual cells. The resulting “barcode_report.tsv” output files, which report the most abundant pair of alpha and beta chains for each cell, were summarized within R. The TRUST4 output was filtered for those cells that were retained in the Seurat analysis and for which the predicted cell type was “abT”. For both the alpha and beta chains, the clonotype was defined as the concatenation of the V and J segments, due to limited calls for the D segment of beta chains. Relative frequencies of clonotypes were computed at both the sample and individual cluster level.

### RNA velocity analysis

Unspliced and splice read counts were computed using velocyto v0.17.17 (La Manno *et al*., 2018). As input for each sample, velocyto (with the “run10x” command) was given the output directory of CellRanger, the CellRanger mouse reference gene annotation (mm10, v3.0.0), and an annotation of repetitive elements for the mouse genome (mm10 RepeatMasker track downloaded from the UCSC Genome Browser in GTF format) (Navarro Gonzalez et al., 2021). The resulting read counts were analyzed with the scVelo v0.2.4 Python package (Bergen et al., 2020). Cells were filtered to those that were analyzed with Seurat and annotated with the Seurat-computed clusters. After standard preprocessing documented by scVelo, velocities were computed using its “stochastic” model. Genes with velocities that were significantly higher in one cluster compared to cells from all other clusters were identified using the “rank_velocity_genes” method (with min_corr =0.3).

### RT-qPCR experiments

CD44^+^Tetramer^+^ cells were harvested from lung and spleen following vaccination and pathogen challenge described above. The lungs were processed, stained, and sorted in the same manner as explained before for scRNAseq. Spleens were mashed through 40 μm filters, and then subjected to red blood cell lysis (ACK buffer, Gibco™, Cat#A1049201) for 3 minutes at room temperature. Samples were washed with 15 mL of wash buffer (RPMI with 1% FBS) and the CD4+ T cells were enriched using MojoSort™ Mouse CD4 cell isolation (Biolegend, Cat#480006). The cells were stained and sorted as explained before in scRNAseq section as well. Both lungs and spleen were sorted in 1.5 ml microtubes with 0.5% BSA in PBS 30 minutes after surface staining and kept at 4°C. Lung and spleen cells were not fixed after surface staining and sorted in sterile condition in order to perform *in vitro* stimulation experiments. The samples were pooled after surface staining step (3-4 mice/sample, total of 3 samples).

For stimulation studies, cells from lung or spleen (50,000-200,000 cells/well) were left unstimulated (RPMI with 0.5% BSA) or stimulated with IFNα (10,000 units/mL) or IFNγ (10 ng/ml) and analyzed by RT-qPCR at 12 hours. To measure mRNA expression levels of genes, cDNA was generated directly from cell lysate using the Invitrogen SuperScript IV CellsDirect cDNA Synthesis Kit (ThermoFisher Scientific, 11750150). qPCR was performed on a Rotor-Gene Q system (Qiagen) using TaqMan Gene Expression Assays (ThermoFisher Scientific, Ifi204 Mm00492602_m1; Mx1 Mm00487796_m1; Pml Mm00476969_m1; Slfn5 Mm00806095_m1; Ifit1 Mm00515153_m1; Ifitm3 Mm00847057_s1; Il17a Mm00439618_m1; 18S 4319413E) and TaqMan Fast Advanced Master Mix (ThermoFisher Scientific, 4444556). Relative quantification was performed by the ΔΔCT method with 18S as a reference gene. Relative expression levels were compared using data from one experiment representative of three independent experiments using two-tailed Student’s t-test.

## Data Availability

Raw and integrated scRNAseq data is deposited in Gene Expression Omnibus (GEO)M database, with accession number GSE198466.

## ACKNOWLEDGMENTS

The authors thank Dr. Jenny Gumperz for her insight and expertise regarding natural killer T cells and the University of Wisconsin-Madison Clinical Cancer Center, Flow Cytometry Core Facility (UWCCC) and Biotechnology Center for help with cell sorting and sequencing. Robert Gordon (Department of Pediatrics) assisted with graphic illustration. The UWCCC is supported by NIH Shared Instrumentation Grant 1S100OD018202–01 and University of Wisconsin Carbone Cancer Center Support grant P30 CA014520. We also acknowledge support from NIH grants R01 AI130411, U01AI124299, R01 AI035681 (to B.K.); R01AI168370 (to B.K. and M.W.); R01 AI093553 (to M.W.); the University of Wisconsin-Madison Institutional Clinical and Translational Science Award UL1 TR002373 (to C.D.); and from the University of Wisconsin School of Medicine and Public Health Pediatrics Fellow/Resident research grant (to T.W.). Xin He is a Cancer Research Institute Irvington Fellow supported by the Cancer Research Institute (CRI4476). Bruce Klein is a CIFAR Fellow and receives support from this agency.

## SUPPLEMENTAL FIGURES

**Supplemental Figure 1. (a)** Flow cytometry plots for selection of tetramer^+^ CD4^+^ T cells from IN sample. **(b)** Counts of sorted and sequenced tetramer-positive CD4^+^ cells from SC and IN samples. **(c)** UMAP for cell cycle phases of IN and SC groups. Cells in all stages are distributed across all clusters. **(d)** UMAP after regression of cell cycle genes in IN and SC groups shows cohesive clusters with both the original cluster assignments prior to regression (top) and the new cluster assignments after regression (bottom). **(e)** Stacked bar plot depicting relative abundance of each cluster within the IN and SC groups.

**Supplemental Figure 2. (a)** Heatmap showing average expression of tissue-associated memory T cell genes involved in cytoskeleton, cell matrix, membrane scaffolding and adhesion. Gene list adapted from Szabo et al.(Szabo *et al*., 2019) The cluster with conspicuous downregulation of these tissue-associated genes corresponds to the naïve/T_CM_ population (cluster 9). **(b)** Comparison of the number of detected genes and reads in quiescent cells (cluster 11) compared to other clusters. Both were significantly lower in cluster 11 cells (genes, P=1.5×10^−174^; reads, P=6.1×10^−103^). P-values generated with Mann-Whitney test (***= P<0.001). **(c)** Less than 5% of reads mapped to mitochondrial genes across all clusters. Cluster 11 cells were comparable to other clusters for percent mitochondrial reads. **(d)** Comparison of the number of detected genes and reads in myeloid-like T cells (MyT, cluster 16) compared to other clusters. There was no significant increase in gene or read counts to suggest doublets within this population (genes, P=0.17; reads, P=0.39). P-values generated with Mann-Whitney test. **(e)** UMAP for Th1 transcription factor *Tbx21* (Tbet) and Th17 transcription factor *Rorc* (RORγt).

**Supplemental Figure 3. (a)** Total cells with α, β, or αβ TCR clonotype calls using TRUST4 in IN and SC samples. **(b)** Proportion of cells within each cluster with an α, β, or αβ TCR clonotype call. **(c)** Frequency of the most abundant α, β, or αβ TCR clonotype was higher IN than SC groups. **(d)** Cumulative frequency of the top α, β, or αβ TCR clonotypes. **(e)** The proportion of the 5 most abundant TCR α-chain clonotypes within each cluster.

**Supplemental Figure 4. (a)** UMAP for expression of T cell anergy gene *Rnf128* (GRAIL), separated by IN and SC samples. **(b)** Violin plot showing lack of expression of innate-like T cell gene *Zbtb16* (PZLF) in NK-like Th1 and all other cells.

**Supplemental Figure 5. (a)** Quantitative reverse transcriptase PCR (RT-qPCR) detects increased expression of Tis T cell marker genes (*Ifi204, Mx1, Pml, Slfn5, Ifit1*) in tetramer-positive, CD44-positive (black) cells compared to control CD44-negative (white) cells from the spleen of *Blastomyces*-challenged, subcutaneously vaccinated mice. The addition of IFNα increased expression of these genes in both tetramer-positive and control cells. **(b)** RT-qPCR for Tis T cell marker genes in tetramer-positive, CD44-positive cells from the spleen of *Blastomyces*-challenged, subcutaneously vaccinated mice do not increase 12h after IFNγ stimulation compared to those left unstimulated (Mock). **(c)** RT-qPCR for *Ifit3* in tetramer-positive, CD44-positive cells from the lungs of *Blastomyces*-challenged, subcutaneously vaccinated mice do not increase 12h after IFNγ stimulation compared to those left unstimulated (Mock). **(d)** Percentage of CD4^+^ T cells and tetramer-positive T cells harvested from the lungs of IFNγR^-/-^ (KO) mice and wild-type mice that were vaccinated SC, experimentally challenged via the pulmonary route, and analyzed three days post-infection.

