## Supplemental data for "Novel populations of CD4^+^ T cells associated with vaccine efficacy"

a.

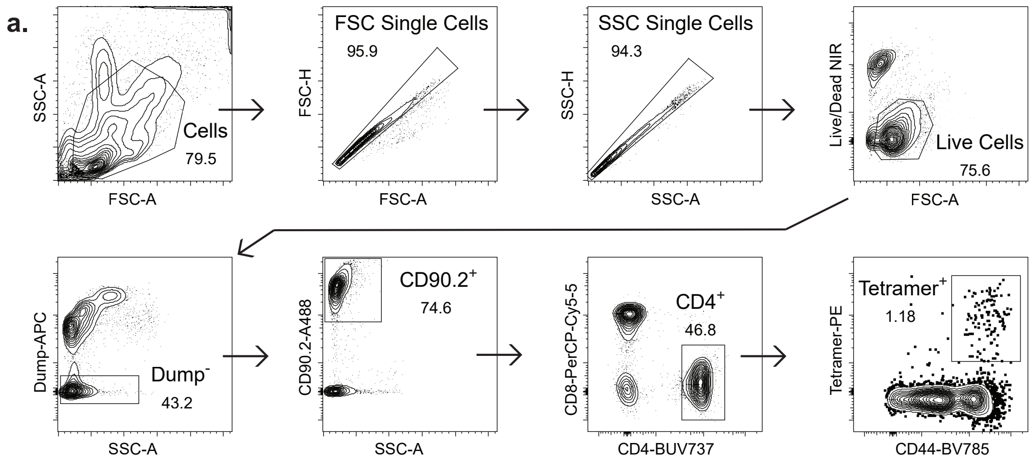


b.

| Count | SC | IN |
|---|---|---|
| Sorted tetramer⁺ cells | 650,000 | 21,000 |
| Sequenced cells | 11,977 | 8,661 |

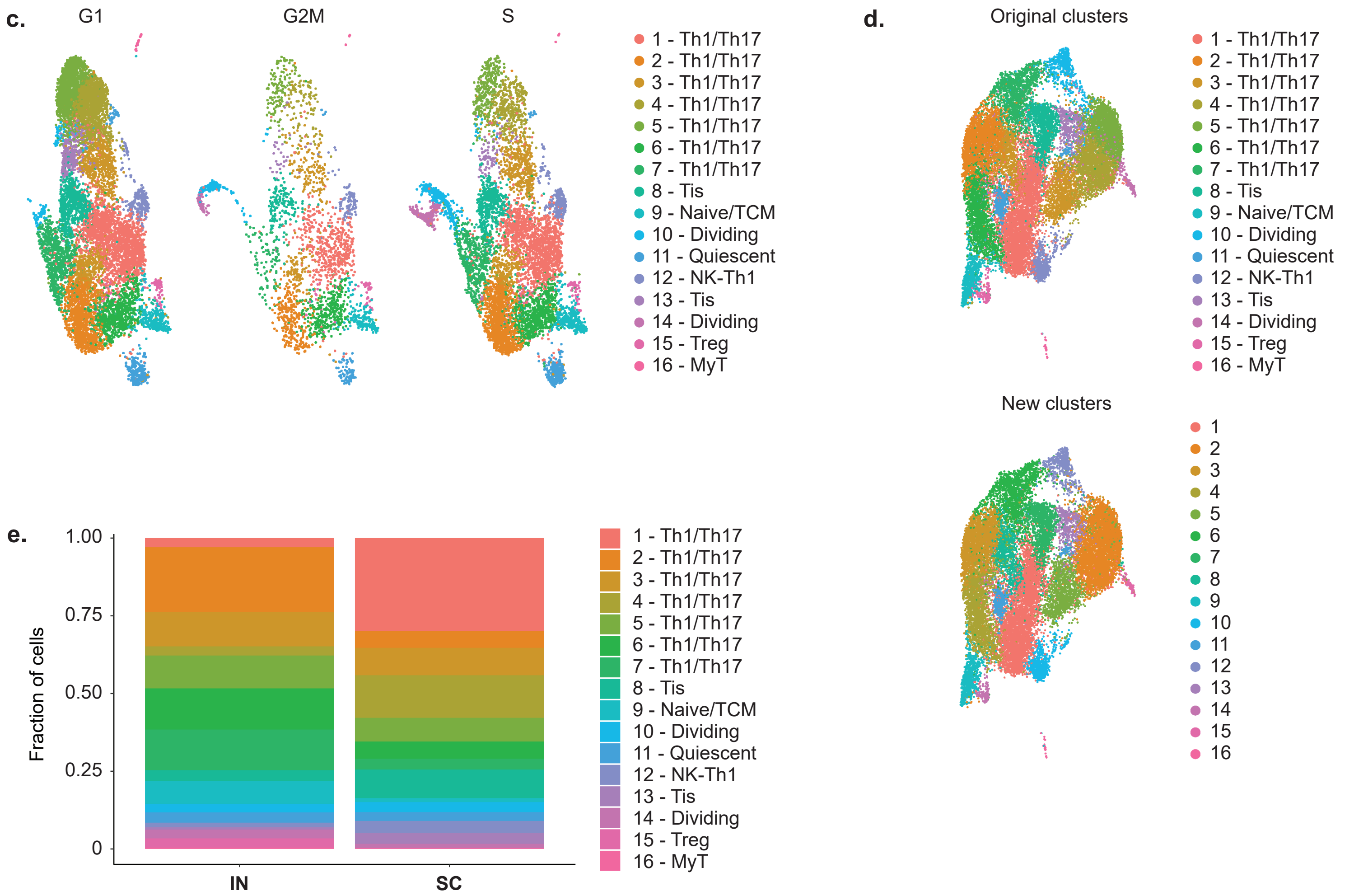

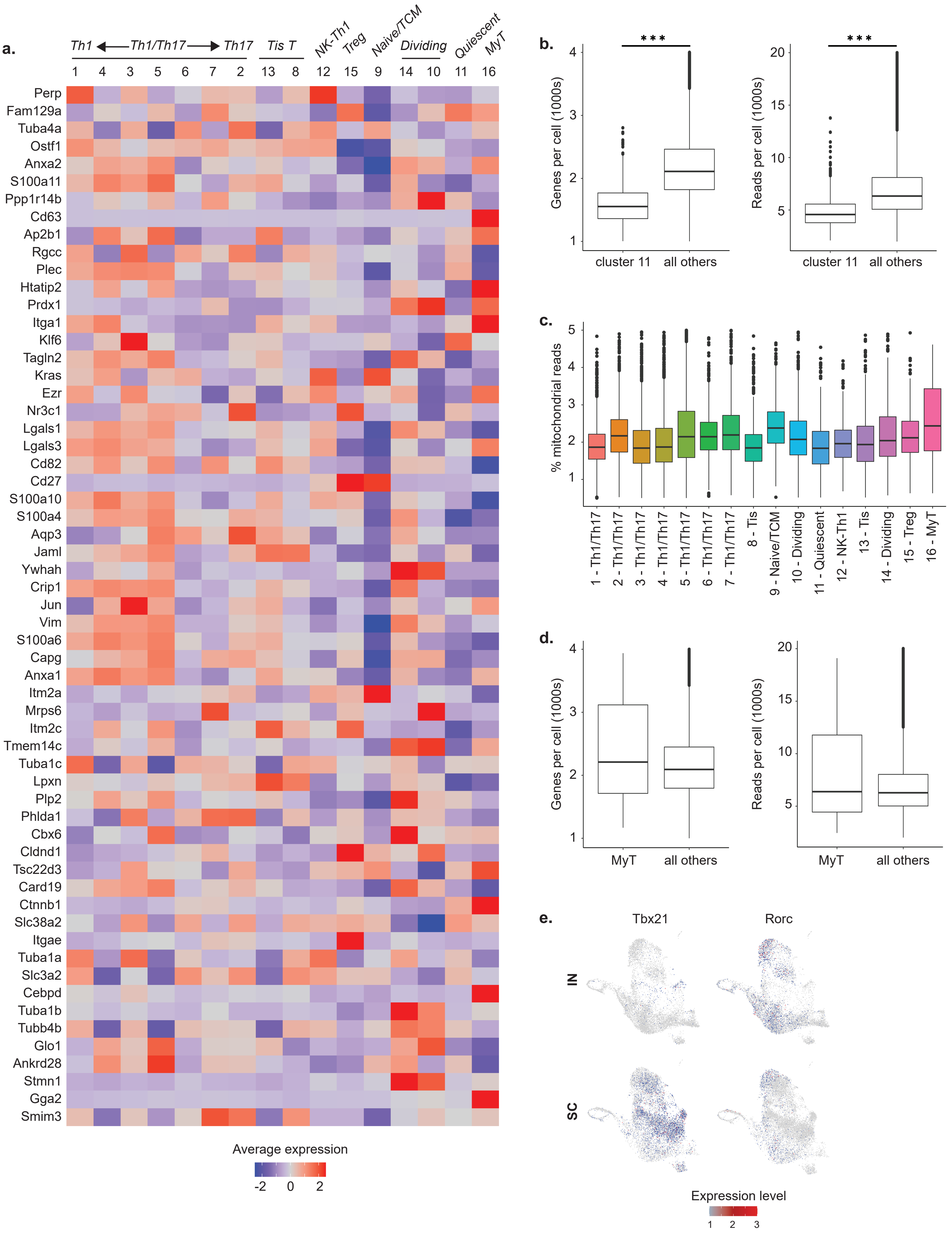
a.
Th1
Th1/Th17
Th17
Tis T
NK-Th1
Treg
Naive/TCM
Dividing
Quiescent
MyT
1 4 3 5 6 7 2 13 8 12 15 9 14 10 11 16
Perp
Fam129a
Tuba4a
Ostf1
Anxa2
S100a11
Ppp1r14b
Cd63
Ap2b1
Rgcc
Plec
Htatip2
Prdx1
Itga1
Klf6
Tagln2
Kras
Ezr
Nr3c1
Lgals1
Lgals3
Cd82
Cd27
S100a10
S100a4
Aqp3
Jaml
Ywhah
Crip1
Jun
Vim
S100a6
Capg
Anxa1
Itm2a
Mrps6
Itm2c
Tmem14c
Tuba1c
Lpxn
Plp2
Phlda1
Cbx6
Cldnd1
Tsc22d3
Card19
Ctnnb1
Slc38a2
Itgae
Tuba1a
Slc3a2
Cebpd
Tuba1b
Tubb4b
Glo1
Ankrd28
Stmn1
Gga2
Smim3
Average expression
-2 0 2
b.
***
Genes per cell (1000s)
1 2 3 4
cluster 11
all others
Reads per cell (1000s)
5 10 15 20
cluster 11
all others
c.
% mitochondrial reads
1 2 3 4 5
1 - Th1/Th17
2 - Th1/Th17
3 - Th1/Th17
4 - Th1/Th17
5 - Th1/Th17
6 - Th1/Th17
7 - Th1/Th17
8 - Tis
9 - Naive/TCM
10 - Dividing
11 - Quiescent
12 - NK-Th1
13 - Tis
14 - Dividing
15 - Treg
16 - MyT
d.
Genes per cell (1000s)
1 2 3 4
MyT
all others
Reads per cell (1000s)
5 10 15 20
MyT
all others
e.
Tbx21
Rorc
IN
SC
Expression level
1 2 3

a.

| Sample | Total αβ T cells analyzed | # cells with α chain call | # cells with β chain call | # cells with both chains called |
|---|---|---|---|---|
| IN | 2421 | 1880 | 735 | 212 |
| SC | 2619 | 1933 | 822 | 173 |

b.

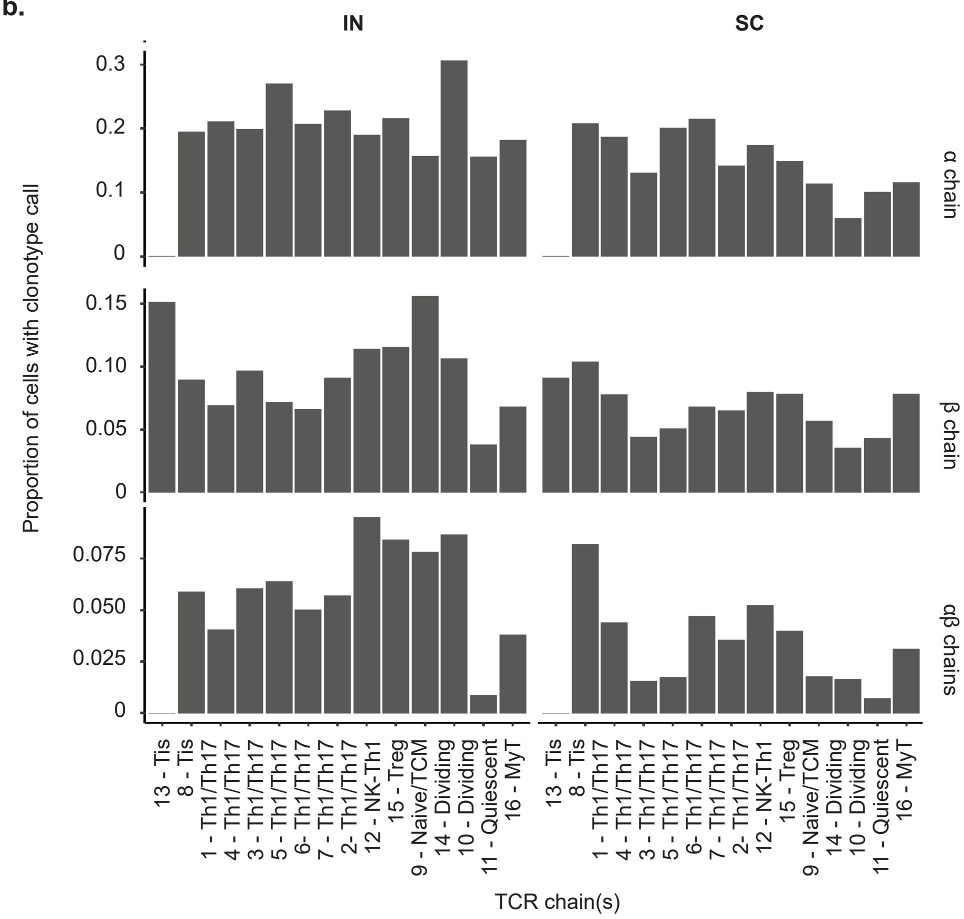


c.

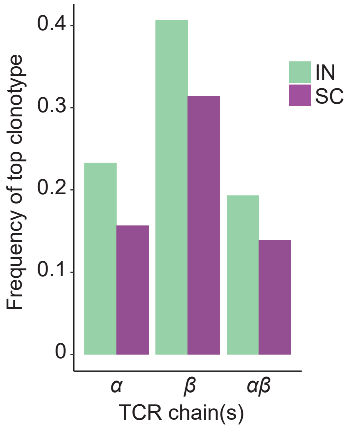


d.

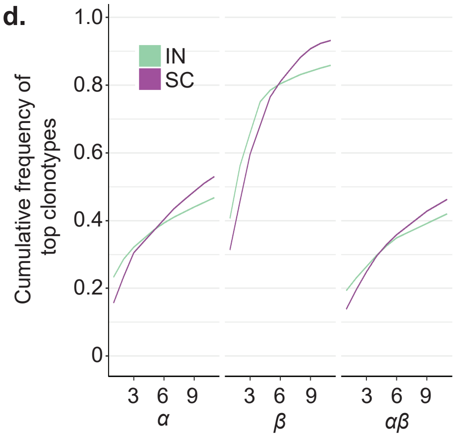


e.

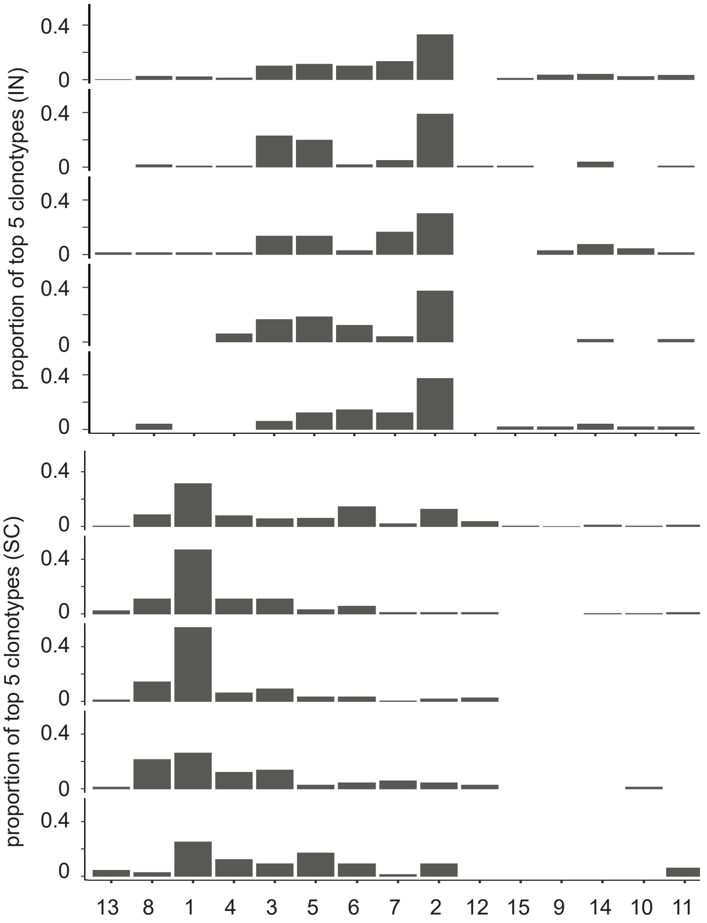

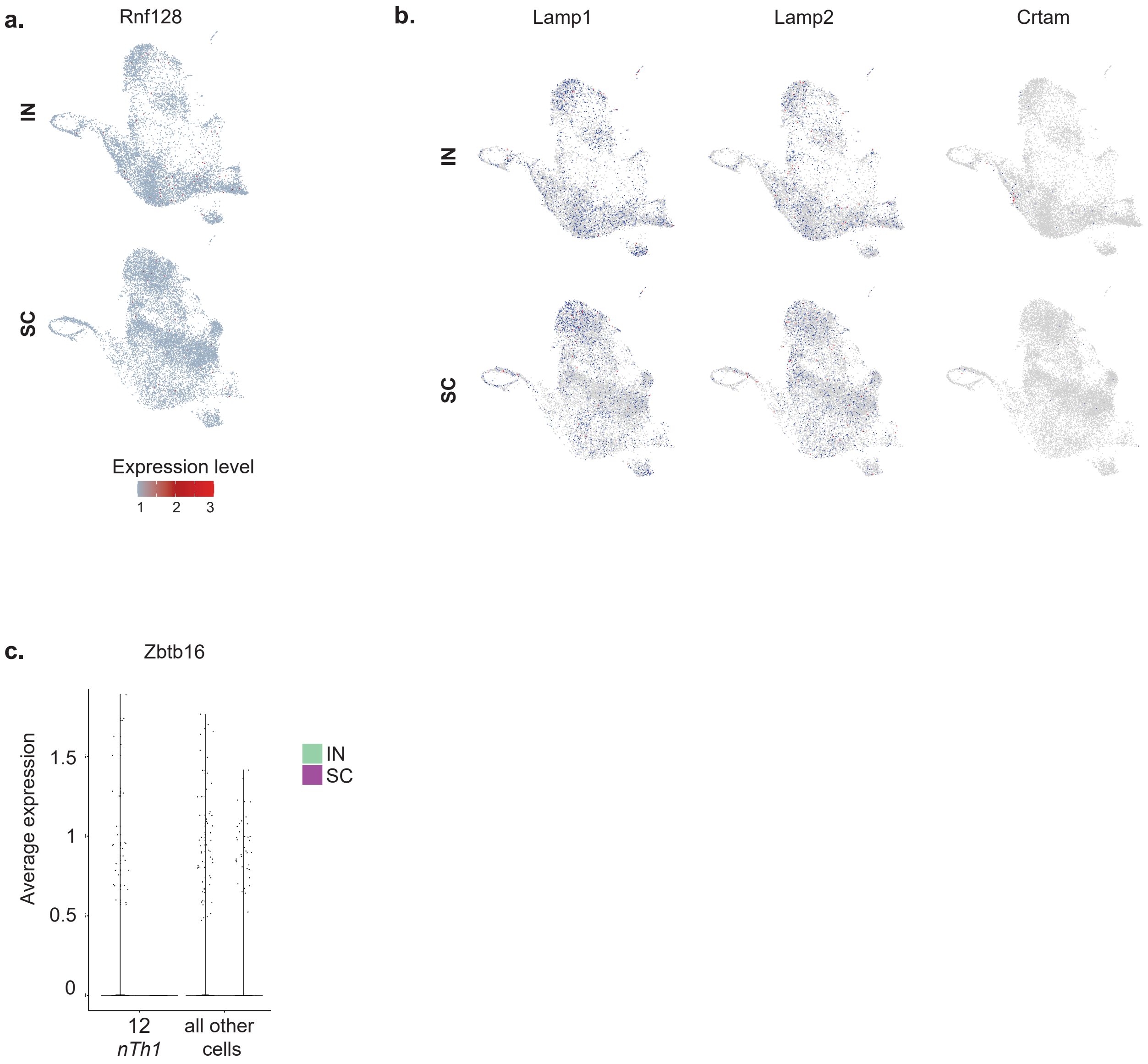
a.
Rnf128
IN
SC
Expression level
1
2
3
b.
Lamp1
Lamp2
Crtam
IN
SC
c.
Zbtb16
Average expression
1.5
1
0.5
0
12
nTh1
all other
cells
IN
SC

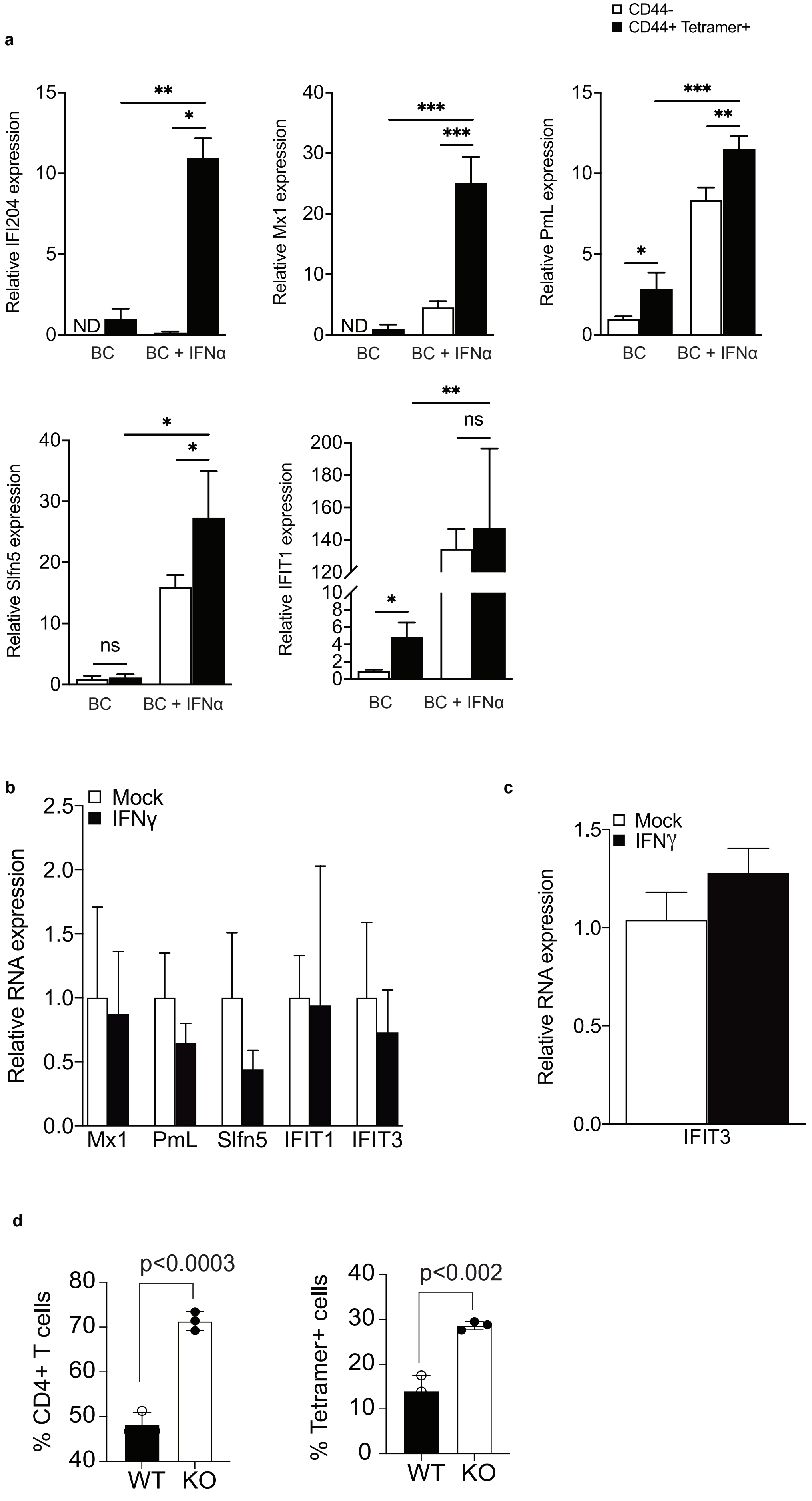

a
CD44-
CD44+ Tetramer+
Relative IFI204 expression
Relative Mx1 expression
Relative PmL expression
Relative Slfn5 expression
Relative IFIT1 expression
ND
ns
BC
BC + IFNα
b
Mock
IFNγ
Relative RNA expression
Mx1
PmL
Slfn5
IFIT1
IFIT3
c
Mock
IFNγ
Relative RNA expression
IFIT3
d
% CD4+ T cells
p<0.0003
% Tetramer+ cells
p<0.002
WT
KO

| Cluster | All cells | Percent (All) | | IN | Percent (IN) | SC | Percent (SC) |
|---|---|---|---|---|---|---|---|
| 1 | 3842 | 18.6% | | 254 | 2.9% | 3588 | 30.0% |
| 2 | 2453 | 11.9% | | 1807 | 20.9% | 646 | 5.4% |
| 3 | 2005 | 9.7% | | 956 | 11.0% | 1049 | 8.8% |
| 4 | 1910 | 9.3% | | 259 | 3.0% | 1651 | 13.8% |
| 5 | 1829 | 8.9% | | 916 | 10.6% | 913 | 7.6% |
| 6 | 1799 | 8.7% | | 1143 | 13.2% | 656 | 5.5% |
| 7 | 1556 | 7.5% | | 1133 | 13.1% | 423 | 3.5% |
| 8 | 1394 | 6.8% | | 301 | 3.5% | 1093 | 9.1% |
| 9 | 795 | 3.9% | | 636 | 7.3% | 159 | 1.3% |
| 10 | 632 | 3.1% | | 242 | 2.8% | 390 | 3.3% |
| 11 | 627 | 3.0% | | 285 | 3.3% | 342 | 2.9% |
| 12 | 577 | 2.8% | | 125 | 1.4% | 452 | 3.8% |
| 13 | 485 | 2.4% | | 65 | 0.8% | 420 | 3.5% |
| 14 | 416 | 2.0% | | 253 | 2.9% | 163 | 1.4% |
| 15 | 274 | 1.3% | | 253 | 2.9% | 21 | 0.2% |
| 16 | 44 | 0.2% | | 33 | 0.4% | 11 | 0.1% |
| **Total** | 20,638 | 100% | | 8661 | 100% | 11,977 | 100% |

**Supplemental Table 1.** Number and percent of cells in each cluster overall (All cells) and in intranasally vaccinated (IN) and subcutaneously vaccinated (SC) mice.

| gene symbol | avg_log2FC | pct.SC | pct.IN | | gene symbol | avg_log2FC | pct.SC | pct.IN |
|---|---|---|---|---|---|---|---|---|
| Il17a | -2.939 | 0.166 | 0.71 | | Nkg7 | 1.944 | 0.668 | 0.118 |
| Il17f | -1.876 | 0.062 | 0.398 | | Rps17 | -0.532 | 0.92 | 0.972 |
| Il18rap | 0.858 | 0.532 | 0.208 | | Ifitm2 | 1.788 | 0.437 | 0.047 |
| Ctla4 | -1.262 | 0.483 | 0.75 | | Ifitm3 | 2.028 | 0.453 | 0.04 |
| Ramp1 | -1.120 | 0.396 | 0.709 | | Ifngr1 | 0.595 | 0.919 | 0.823 |
| Rgs1 | -0.875 | 0.908 | 0.983 | | Rps12 | -0.406 | 0.998 | 0.999 |
| Rgs16 | -1.117 | 0.217 | 0.472 | | Lilr4b | 0.859 | 0.429 | 0.175 |
| Vim | 0.685 | 0.988 | 0.939 | | Rps15 | -0.640 | 0.981 | 0.996 |
| Rpl12 | -0.549 | 0.995 | 0.998 | | Gadd45b | -0.961 | 0.745 | 0.88 |
| Itga4 | -1.062 | 0.246 | 0.659 | | Ifng | 2.193 | 0.63 | 0.142 |
| AA467197 | 1.762 | 0.73 | 0.164 | | Rps26 | -0.449 | 0.996 | 0.998 |
| Atp8b4 | 0.707 | 0.345 | 0.04 | | Dusp4 | -1.012 | 0.118 | 0.496 |
| Dusp2 | 1.303 | 0.77 | 0.604 | | Uba52 | -0.361 | 0.982 | 0.996 |
| 1110034G24Rik | -0.528 | 0.065 | 0.277 | | Junb | -0.632 | 0.979 | 0.996 |
| Pmepa1 | -0.653 | 0.194 | 0.471 | | Znrf1 | -0.665 | 0.605 | 0.817 |
| Ar | -0.551 | 0.153 | 0.386 | | Gabarapl2 | 0.576 | 0.898 | 0.793 |
| Car5b | 0.579 | 0.319 | 0.077 | | Itgb1 | 1.457 | 0.909 | 0.582 |
| Sema4a | 0.594 | 0.374 | 0.113 | | Lgals3 | 1.996 | 0.81 | 0.284 |
| S100a13 | 0.572 | 0.916 | 0.823 | | Dnajc15 | 1.188 | 0.799 | 0.427 |
| S100a6 | 0.915 | 0.973 | 0.86 | | Epsti1 | 1.277 | 0.759 | 0.345 |
| S100a10 | 0.787 | 0.992 | 0.966 | | Rps25 | -0.362 | 0.99 | 0.997 |
| Cd2 | 0.645 | 0.889 | 0.753 | | Nptn | 0.740 | 0.65 | 0.377 |
| Csf1 | 0.690 | 0.266 | 0.035 | | Rplp1 | -0.506 | 0.999 | 1 |
| Rps20 | -0.423 | 0.998 | 0.999 | | Nt5e | -1.121 | 0.155 | 0.571 |
| Tox | -0.772 | 0.058 | 0.375 | | Chst2 | -0.586 | 0.098 | 0.343 |
| Rps8 | -0.308 | 0.999 | 1 | | Gpx1 | -0.856 | 0.62 | 0.834 |
| Sh3bgrl3 | 0.678 | 0.993 | 0.972 | | Ccr4 | -0.547 | 0.053 | 0.284 |
| Rcc2 | -0.676 | 0.412 | 0.644 | | Cxcr6 | 0.701 | 0.881 | 0.674 |
| Fosl2 | -0.784 | 0.41 | 0.655 | | Ccr2 | 0.900 | 0.69 | 0.327 |
| Arap2 | 0.626 | 0.66 | 0.435 | | Ramp3 | -0.708 | 0.11 | 0.354 |
| Rpl5 | -0.548 | 0.989 | 0.997 | | Hs3st3b1 | 0.530 | 0.294 | 0.083 |
| Selplg | 0.604 | 0.876 | 0.736 | | Rpl23a | -0.461 | 0.969 | 0.988 |
| Ndufa4 | -0.893 | 0.468 | 0.801 | | Ccl5 | 2.212 | 0.327 | 0.071 |
| 1810058I24Rik | 0.697 | 0.709 | 0.5 | | Hlf | -0.931 | 0.113 | 0.45 |
| Zyx | 0.768 | 0.581 | 0.306 | | Tbx21 | 0.841 | 0.397 | 0.052 |
| Arl6ip5 | 0.724 | 0.842 | 0.662 | | Ikzf3 | -1.184 | 0.153 | 0.597 |
| Slc2a3 | 0.727 | 0.488 | 0.198 | | Gna13 | -0.812 | 0.769 | 0.912 |
| Klrd1 | 1.109 | 0.339 | 0.04 | | Timp2 | -0.473 | 0.103 | 0.319 |
| Klrk1 | 1.137 | 0.552 | 0.15 | | Serpinb9 | 0.692 | 0.313 | 0.097 |
| Klrc2 | 0.842 | 0.313 | 0.016 | | Odc1 | -0.732 | 0.811 | 0.935 |
| Klrc1 | 1.344 | 0.479 | 0.026 | | Ahr | -0.605 | 0.104 | 0.354 |
| Ybx3 | 0.663 | 0.71 | 0.487 | | Tspan13 | -0.813 | 0.255 | 0.588 |
| Kcnn4 | 0.662 | 0.694 | 0.471 | | Rps29 | -0.335 | 1 | 1 |
| Zfp36 | -1.032 | 0.499 | 0.76 | | Zfp36l1 | -0.753 | 0.758 | 0.893 |
| Fxyd5 | 0.584 | 0.976 | 0.949 | | Fos | -0.864 | 0.567 | 0.812 |

**Supplemental Table 2**. Top differentially expressed genes in subcutaneously inoculated mice (SC) compared to intranasally inoculated mice (IN). All differences in expression are statistically significant ($P<0.01$). avg_log2FC = average log2 fold change, where positive denotes genes upregulated SC. Ptc.SC = percent of cells expressing gene SC. Pct.IN = percent of cells expressing gene IN.

| gene symbol | avg_log2FC | pct.Tis | pct.other | | gene symbol | avg_log2FC | pct.Tis | pct.other |
|---|---|---|---|---|---|---|---|---|
| Stat1 | 1.463588 | 0.874 | 0.43 | | Dhx58 | 0.552192 | 0.271 | 0.034 |
| Sp100 | 1.021642 | 0.845 | 0.538 | | Ifi35 | 0.956346 | 0.698 | 0.314 |
| Ifi206 | 1.364668 | 0.678 | 0.143 | | Lgals3bp | 1.005164 | 0.608 | 0.206 |
| Ifi214 | 1.158886 | 0.49 | 0.057 | | Rnf213 | 1.640465 | 0.742 | 0.163 |
| Ifi213 | 1.474788 | 0.673 | 0.102 | | Rsad2 | 1.154667 | 0.393 | 0.025 |
| Ifi209 | 1.280381 | 0.672 | 0.194 | | Cmpk2 | 1.025174 | 0.437 | 0.046 |
| Ifi208 | 1.29585 | 0.605 | 0.12 | | Ly6a | 1.384124 | 0.957 | 0.654 |
| Ifi204 | 1.342273 | 0.426 | 0.025 | | Rtp4 | 1.447903 | 0.635 | 0.068 |
| Mndal | 1.548596 | 0.935 | 0.601 | | Parp14 | 1.16424 | 0.624 | 0.203 |
| Ifi211 | 0.920783 | 0.309 | 0.043 | | Dtx3l | 0.876607 | 0.518 | 0.146 |
| Ifi203 | 1.643232 | 0.941 | 0.567 | | Parp9 | 0.847324 | 0.507 | 0.13 |
| Ifih1 | 0.869165 | 0.407 | 0.061 | | Mx1 | 0.997256 | 0.318 | 0.023 |
| B2m | 0.671886 | 1 | 0.999 | | H2-T23 | 0.947757 | 0.987 | 0.902 |
| Samhd1 | 1.457596 | 0.89 | 0.557 | | Eif2ak2 | 0.607137 | 0.297 | 0.054 |
| Zbp1 | 1.701802 | 0.831 | 0.235 | | AW112010 | 1.148043 | 0.988 | 0.867 |
| Gbp7 | 1.141752 | 0.65 | 0.229 | | Ms4a4c | 1.125778 | 0.44 | 0.099 |
| Gbp2 | 1.330516 | 0.529 | 0.141 | | Ms4a4b | 1.308764 | 0.972 | 0.758 |
| Ddx58 | 0.906377 | 0.54 | 0.158 | | Ms4a6b | 1.070097 | 0.972 | 0.789 |
| Isg15 | 2.651938 | 0.891 | 0.252 | | Ifit3 | 1.700427 | 0.434 | 0.02 |
| Gbp9 | 0.653588 | 0.332 | 0.07 | | Ifit1 | 1.757272 | 0.53 | 0.032 |
| Gbp6 | 0.927166 | 0.502 | 0.124 | | Psmb8 | 0.797364 | 0.98 | 0.893 |
| Oasl2 | 0.964412 | 0.379 | 0.028 | | Stat2 | 0.674791 | 0.383 | 0.098 |
| Oas3 | 0.925822 | 0.461 | 0.058 | | Ube2l6 | 0.770356 | 0.356 | 0.089 |
| Trafd1 | 1.021619 | 0.578 | 0.18 | | Psmb10 | 0.93669 | 0.79 | 0.461 |
| Samd9l | 1.105671 | 0.72 | 0.339 | | Slfn2 | 0.957202 | 0.92 | 0.696 |
| Herc6 | 1.191133 | 0.596 | 0.137 | | Irf9 | 0.744718 | 0.542 | 0.201 |
| Usp18 | 0.842146 | 0.324 | 0.017 | | Daxx | 0.686932 | 0.388 | 0.109 |
| Isg20 | 1.766755 | 0.779 | 0.219 | | Psme1 | 0.798974 | 0.944 | 0.782 |
| Trim30a | 1.47597 | 0.761 | 0.21 | | Gbp4 | 0.901133 | 0.565 | 0.226 |
| Trim30d | 0.584551 | 0.266 | 0.04 | | H2-T22 | 0.911738 | 0.836 | 0.586 |
| Ifitm3 | 2.196072 | 0.63 | 0.245 | | H2-K1 | 0.49828 | 1 | 0.999 |
| Irf7 | 1.619405 | 0.811 | 0.229 | | 9930111J21Rik2 | 0.925375 | 0.751 | 0.42 |
| Ddx60 | 0.655933 | 0.3 | 0.043 | | Tapbp | 0.888208 | 0.798 | 0.541 |
| Bst2 | 2.361482 | 0.96 | 0.521 | | Parp10 | 0.597853 | 0.401 | 0.125 |
| Phf11b | 1.275133 | 0.654 | 0.219 | | Helz2 | 0.610078 | 0.371 | 0.11 |
| Epsti1 | 1.175369 | 0.898 | 0.554 | | Chmp4b | 0.756142 | 0.923 | 0.777 |
| Pml | 0.653543 | 0.36 | 0.083 | | Psme2 | 0.722376 | 0.897 | 0.714 |
| Shisa5 | 0.872388 | 0.994 | 0.942 | | Sp110 | 0.807667 | 0.626 | 0.325 |
| Irgm1 | 1.015191 | 0.607 | 0.177 | | H2-Q4 | 0.668293 | 0.966 | 0.844 |
| Ifi47 | 1.28474 | 0.726 | 0.266 | | Trim12c | 0.632304 | 0.373 | 0.123 |
| Igtp | 1.217419 | 0.73 | 0.277 | | Trim12a | 0.723719 | 0.563 | 0.26 |
| Xaf1 | 1.258989 | 0.65 | 0.125 | | H2-D1 | 0.394261 | 1 | 1 |
| Slfn5 | 2.008012 | 0.675 | 0.136 | | Ogfr | 0.639643 | 0.544 | 0.257 |
| Slfn8 | 0.983883 | 0.494 | 0.111 | | Tap1 | 0.693251 | 0.741 | 0.472 |
| Slfn1 | 1.470985 | 0.75 | 0.227 | | Nampt | 0.518962 | 0.348 | 0.118 |

**Supplemental Table 3**. Top differentially expressed genes between Tis T cells (combined cluster 8 and 13) and all other cells. All differences in expression are statistically significant ($P<0.01$). avg_log2FC = average log2 fold change, where positive denotes genes upregulated in Tis T cells compared to other cells. Ptc.Tis = percent of cells in clusters 8 & 13 expressing gene. Pct.other = percent of cells in remaining clusters expressing gene.

| gene symbol | avg_log2FC | pct.Tis.8 | pct.Tis.13 | | gene symbol | avg_log2FC | pct.Tis.8 | pct.Tis.13 |
|---|---|---|---|---|---|---|---|---|
| H3f3b | 1.736125 | 1 | 0.992 | | Sqstm1 | 1.104345 | 0.808 | 0.381 |
| Junb | 1.946537 | 1 | 0.948 | | Nfkbid | 1.15721 | 0.666 | 0.208 |
| Vps37b | 2.251496 | 0.939 | 0.324 | | Irf2bp2 | 0.955399 | 0.876 | 0.6 |
| Bhlhe40 | 2.005815 | 0.981 | 0.79 | | Ptp4a1 | 1.018083 | 0.758 | 0.338 |
| Nfkbia | 2.075163 | 0.995 | 0.808 | | Ctla4 | 1.332767 | 0.7 | 0.268 |
| Pim1 | 1.994649 | 0.986 | 0.74 | | Srgn | 0.722013 | 0.999 | 0.973 |
| Tnfaip3 | 1.977649 | 0.958 | 0.48 | | Coq10b | 0.929079 | 0.793 | 0.359 |
| Odc1 | 2.346535 | 0.963 | 0.555 | | Cxcr4 | 1.266615 | 0.639 | 0.198 |
| Pnrc1 | 1.484511 | 0.981 | 0.68 | | Hif1a | 0.998846 | 0.913 | 0.612 |
| Csrnp1 | 1.757204 | 0.877 | 0.247 | | Hilpda | 1.225128 | 0.496 | 0.072 |
| Ubc | 1.303885 | 0.999 | 0.899 | | Rrad | 1.24763 | 0.679 | 0.254 |
| Kdm6b | 1.875581 | 0.902 | 0.328 | | Stk17b | 0.821825 | 0.959 | 0.81 |
| Ifrd1 | 1.673152 | 0.918 | 0.439 | | Bcl2a1d | 1.063756 | 0.841 | 0.544 |
| Dusp5 | 1.559009 | 0.928 | 0.528 | | Gadd45b | 1.333962 | 0.91 | 0.637 |
| Btg1 | 1.071251 | 1 | 0.986 | | Nfkbiz | 1.015138 | 0.653 | 0.235 |
| Cdkn1a | 1.759169 | 0.83 | 0.272 | | Dnajb9 | 0.948277 | 0.613 | 0.186 |
| Rgs2 | 1.845596 | 0.925 | 0.526 | | Zfp622 | 0.872833 | 0.728 | 0.313 |
| Nr4a1 | 1.766735 | 0.804 | 0.266 | | Dennd4a | 1.057691 | 0.791 | 0.41 |
| Zc3h12a | 1.443751 | 0.758 | 0.192 | | Sub1 | 0.718955 | 0.976 | 0.864 |
| Tgif1 | 1.413129 | 0.714 | 0.144 | | Wsb1 | 0.844474 | 0.742 | 0.33 |
| Bcl2a1b | 1.476229 | 0.938 | 0.658 | | Fth1 | 0.624196 | 1 | 0.994 |
| Sh2d2a | 1.393376 | 0.914 | 0.61 | | Gpr183 | 1.041411 | 0.803 | 0.476 |
| Ubald2 | 1.533704 | 0.831 | 0.357 | | Tnfrsf4 | 1.420604 | 0.605 | 0.221 |
| Ier5 | 1.233939 | 0.963 | 0.616 | | Atf3 | 1.224609 | 0.627 | 0.233 |
| Srsf5 | 1.00658 | 0.981 | 0.771 | | Ly6a | -0.83014 | 0.949 | 0.981 |
| Gna13 | 1.362324 | 0.912 | 0.548 | | Fosl2 | 1.00056 | 0.533 | 0.142 |
| Hspa5 | 1.019739 | 0.993 | 0.891 | | Sdhaf1 | 0.989205 | 0.587 | 0.19 |
| Eif1 | 0.612944 | 1 | 0.994 | | Dusp2 | 1.372155 | 0.859 | 0.604 |
| Rgs1 | 1.595625 | 0.961 | 0.808 | | Mknk2 | 0.806657 | 0.808 | 0.449 |
| Ddx5 | 0.676566 | 1 | 0.981 | | Skil | 0.77867 | 0.933 | 0.672 |
| Tob1 | 1.164373 | 0.795 | 0.278 | | Rinl | -0.87748 | 0.573 | 0.802 |
| Smad7 | 1.081028 | 0.95 | 0.693 | | Sertad1 | 0.808728 | 0.652 | 0.247 |
| Cish | 1.082261 | 0.918 | 0.614 | | Gramd3 | 1.019667 | 0.732 | 0.379 |
| Socs3 | 1.362927 | 0.707 | 0.212 | | Birc2 | 0.801778 | 0.807 | 0.478 |
| Prdx6 | 1.057144 | 0.938 | 0.711 | | Lmnb1 | 0.927639 | 0.734 | 0.357 |
| Zfp36l2 | 1.328891 | 0.941 | 0.691 | | Arl6ip1 | -0.7051 | 0.863 | 0.942 |
| Hist1h1c | 1.982579 | 0.73 | 0.318 | | Tubb4b | 0.795374 | 0.83 | 0.507 |
| Rgcc | 1.262302 | 0.896 | 0.538 | | Ube2s | 0.714336 | 0.881 | 0.623 |
| Ier2 | 1.064392 | 0.944 | 0.676 | | Rora | 0.850761 | 0.919 | 0.765 |
| Slc3a2 | 1.048187 | 0.877 | 0.53 | | Ppp1r15a | 0.802229 | 0.863 | 0.52 |
| Nr4a3 | 1.165843 | 0.628 | 0.151 | | Arid5a | 0.834068 | 0.702 | 0.322 |
| Gpr132 | 1.059531 | 0.828 | 0.431 | | Vgll4 | 0.802229 | 0.761 | 0.406 |
| Arf4 | 0.89394 | 0.966 | 0.736 | | Pde4b | 0.911252 | 0.709 | 0.379 |
| Per1 | 1.036051 | 0.613 | 0.146 | | Sytl3 | 0.881117 | 0.788 | 0.458 |
| Zfp36l1 | 1.050164 | 0.905 | 0.588 | | Ccnl1 | 0.774097 | 0.89 | 0.635 |

**Supplemental Table 4**. Top differentially expressed genes between cluster 8 and cluster 13 Tis T cells. All differences in expression are statistically significant (P<0.01). avg_log2FC = average log2 fold change, where positive denotes genes upregulated in cluster 8 compared to cluster 13. Ptc.Tis.8 = percent of cells in cluster 8 expressing gene. Pct.Tis.13 = percent of cells in cluster 13 expressing gene.

| Cluster 8 Tis T cells | Cluster 13 Tis T cells |
|---|---|
| Ifi213 | Epsti1 |
| Stat2 | Sp100 |
| Parp14 | Nxpe3 |
| Il18rap | Rnf213 |
| Serpinb9 | Smyd3 |
| Gm4070 | Ddx60 |
| Slfn1 | Slco3a1 |
| Tnfrsf4 | Herc6 |
| Ddx60 | Skap1 |
| Ptp4a2 | Samhd1 |

**Supplemental Table 5**. Genes with the most significantly different velocities in Tis T cells, identified from analysis of unspliced reads with Velocyto.[1] The only gene that also appears among the top differentially expressed genes between cluster 8 and 13 is *Tnfrsf4*, which is already more highly expressed in cluster 8. Data shown for Tis T cell-enriched SC sample only.

1. La Manno G, Soldatov R, Zeisel A, et al. RNA velocity of single cells. *Nature*. 08 2018;560(7719):494-498. doi:10.1038/s41586-018-0414-6
